# New HisKA-like HMM profiles uncover insights on histidine kinase diversity

**DOI:** 10.64898/2026.02.27.708494

**Authors:** Louison Silly, Guy Perrière, Philippe Ortet

## Abstract

**Background:** Histidine kinases (HKs) are key components of many signaling pathways, particularly through their involvement in two-component systems (TCS). By utilizing autophosphorylation and phosphotransfer to response regulators (RRs), they enable organisms to adapt to their environment. Most HKs are transmembrane proteins featuring a sensing domain located outside the cell and two catalytic domains, HisKA and HATPase. The HATPase domain mediates interaction with ATP, while HisKA harbors the phosphorylatable histidine residue. HKs participate in a wide range of environmental adaptation mechanisms, including light sensing and responses to biochemical changes. Characterizing their diversity is therefore essential to better understand how cells interact with their environment. Incomplete HKs (iHKs), lacking either the HisKA or HATPase domain, have been described. Among iHKs that retain an HATPase domain, some possess a region of their sequence where an HisKA domain would be expected. These iHKs may include “true” HKs harboring novel, uncharacterized HisKA domains, which could fill gaps in various signaling pathways.

**Results:** In this study, we analyzed 869 964 iHK sequences carrying an HATPase domain but lacking an HisKA domain. We generated 18 HisKA-like HMM profiles and conducted multiple meta-analyses to assess their HisKA-like characteristics. Their predicted 3D structures were found to match those of known HisKA domains. Furthermore, the genomic context of genes associated with these profiles revealed the presence of genes implicated in signal transduction pathways. Several profiles were cross-validated against curated annotations, as well as against a “negative dataset” composed of non-HK proteins. Lastly, using these profiles we identified 112 iHKs across 22 model organisms of known interest.

**Conclusions:** This work describes 18 HisKA-like profiles identified in prokaryotic sequences, supported by multiple lines of evidence for their HisKA characteristics. These profiles, along with the complete methodology used to identify them, are available in the data repository https://doi.org/10.57745/Y9NOP9 and the git repository https://gitlab.in2p3.fr/louison.silly/hiska-like-domain-characterization. We encourage the integration of these profiles into genome annotation pipelines, as we believe they could facilitate the identification of novel HKs in prokaryotic regulatory pathways, both in model and non-model organisms.

## 1 Introduction

Microorganisms are subject to a wide range of environmental stimuli, from light intensity to physical and biochemical changes (Gomelsky and Hoff, 2011, O’Toole and Wong, 2016, Krämer, 2010, Matilla et al., 2022). In the microbial world, one common mechanism of adaptation to these stimuli is signal transduction through protein phosphorylation. Notable examples include serine/threonine protein kinases (STKs) and histidine kinases (HKs), the latter being particularly prevalent in prokaryotes. HKs are the first component of two-component systems (TCSs), found in most bacteria. TCSs are also present in archaea (Gumerov et al., 2023, Barakat et al., 2011) and in some eukaryotes (Koretke et al., 2000), unlike one-component systems, which are specific to prokaryotes.

Classical TCSs comprise two proteins: an HK and a response regulator (RR) (Stock et al., 1999). Most HKs are transmembrane proteins with an input domain located outside the cell. Upon sensing a signal through this domain, an HK undergoes autophosphorylation within a homodimer. The phosphate group is subsequently transferred to the HK’s cognate RR on a specific aspartate residue within a receiver (REC) domain, ultimately resulting in the regulation of gene expression. Some TCSs operate as multi-step phosphorelay (MSP) systems, characterized by the addition of a third protein bearing a histidine phosphotransfer (HPT) domain, which acts as an intermediary between the HK and the RR. In most MSPs, the HK exhibits a distinct domain architecture compared to those found in classical TCSs. Frequently, the HK harbors a REC domain and is accordingly referred to as a hybrid histidine kinase (HHK) (Stock et al., 1999, Barakat et al., 2011).

Given their crucial role in signal transduction, HKs have been extensively studied. They can be classified in different ways, based on sequence similarities or domain architectures (Dutta et al., 1999, Wuichet and Zhulin, 2010, Barakat et al., 2011), and both approaches are employed in databases such as MIST and P2CS (Gumerov et al., 2023, Ortet et al., 2015). HKs possess two catalytic domains: HisKA, which harbors the phosphorylatable histidine residue, and HATPase, which mediates interaction with ATP. These domains are described in databases such as Pfam (Paysan-Lafosse et al., 2025), SMART (Letunic et al., 2021), and PROSITE (Sigrist et al., 2025), either as two distinct profiles or as a single combined profile. In Pfam and SMART, for instance, HisKA and HATPase are described separately: entries PF00512, PF06580, PF07536, PF07568, PF07730, and PF19191 correspond to various HisKA domains, while entries PF02518, PF13581, PF13589, PF13749, and PF14501 correspond to HATPase domains. Some entries are not annotated as HisKA domains but share structural similarities with them, for example, PF14689, which corresponds to the α-helical domain of the Spo0B protein. In PROSITE, by contrast, a single profile (PS50109) represents both the HisKA and HATPase domains. Despite this extensive catalog of HisKA domains across databases, some proteins with an HK-like domain architecture lack a recognizable HisKA domain. These proteins are referred to as incomplete HKs (iHKs). Some iHKs may represent “true” HKs harboring novel, uncharacterized HisKA domains, and could therefore fill gaps in various signaling pathways. Characterizing their diversity may thus shed light on how cells – particularly in prokaryotes – interact with their environment. In this study, we analyzed more than 800 000 iHK sequences and characterized eighteen novel HisKA-like profile families.

## 2 Methods

The pipeline presented here is schematized in Figure 1.

**Figure 1:**
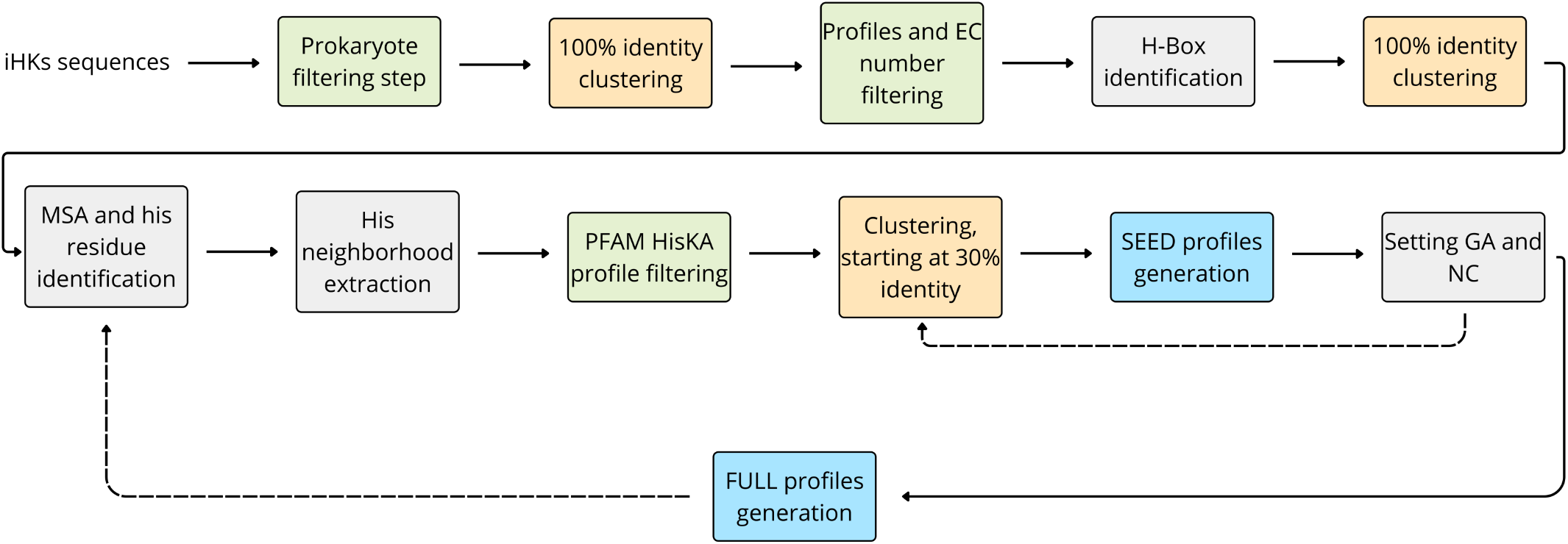
Pipeline used to identify HisKA-like profiles. Filtering steps are highlighted in green, clustering steps in yellow, and profile generation steps in blue.

### 2.1 Sequence retrieval and filtering

Sequences analyzed in this study were taken from an unreleased version of P2CS and are available for download at the link provided in section 6.3. These sequences were initially retrieved from the February 2025 release of RefSeq (Goldfarb et al., 2025) and subsequently annotated as described in (Barakat et al., 2011) to identify HKs and iHKs. We selected sequences lacking an HisKA domain but carrying an HATPase domain, one of the two core domains of HKs, as well as at least one additional domain (most commonly a sensor domain at the N-terminal end). Sequences were divided by taxonomy (bacterial, archaeal, plant, and fungal) and then clustered at 100% identity and coverage using MMseqs2 (Steinegger and Söding, 2018) to remove redundancy. Prior to clustering, bacterial and archaeal sequences were filtered using GTDB v220.0 annotations (Parks et al., 2022) to retain only sequences from high-quality genomes. Completeness and contamination thresholds from CheckM (Parks et al., 2015) and CheckM2 (Chklovski et al., 2023) were set to ≥ 98% and ≤ 1% respectively. All subsequent analyses were performed on the representative sequences of each MMseqs2 cluster.

To minimize false positives and avoid the rediscovery of known domains, multiple filtering steps were implemented. All sequences were annotated using InterProScan (Jones et al., 2014) against the following databases: Pfam v37.0, SMART v9.0, PROSITE v2023_05, SuperFamily v1.75 (Pandurangan et al., 2019), CDD v3.21 (Wang et al., 2023), and CATH-Gene3D v4.4 (Lewis et al., 2018, Sillitoe et al., 2021). EC numbers were assigned to sequences using eggNOG-mapper v2 and the eggNOG database v5.0 (Huerta-Cepas et al., 2019, Cantalapiedra et al., 2021). Since InterProScan relies on FULL alignments, proteins were also aligned using SEED alignments with HMMER v3.4 (Eddy, 2011). Here, FULL and SEED alignments follow the Pfam definitions of these terms: the SEED profile is built from a multiple sequence alignment (MSA) of the initial sequences in which the profile is identified, while the FULL profile is built iteratively by incorporating additional sequences as the profile is identified in new ones. Annotation results were parsed to exclude proteins matching the domains listed in Supplementary Table S1 except for PROSITE profile PS50109, or assigned an EC number not listed in Supplementary Table S2.

We then searched for matches with profile PS50109, which covers both HisKA and HATPase domains in PROSITE, in contrast to the separate HisKA and HATPase profiles used in Pfam. Since iHKs matching this profile lack any known HisKA domain by definition, such matches are likely attributable to their HATPase domain. Nevertheless, a match with PS50109 increases the likelihood that these sequences represent “true” HKs, as they would be annotated as such under the PROSITE framework.

### 2.2 Identification of H-Box and conserved histidine

The H-Box refers to the region of an HK containing the phosphorylatable histidine residue. We searched for H-Boxes in the region immediately upstream of the HATPase domain in our iHKs. The search window was defined relative to the HATPase domain, spanning from −130 to −30 amino acids upstream of the domain boundary, and was required to begin at a position ≥ 75 relative to the start of the sequence, so as to avoid searching for a potential HisKA domain within the N-terminal region. Other annotations (e.g. domains identified by the P2CS pipeline) were also taken into account to prevent searching within already annotated domains. Identified H-Boxes were clustered at 100% identity and coverage, and representatives were then merged across taxonomic groups.

H-Box sequences were aligned using MAFFT v7 (Katoh and Standley, 2013) with the L-INS-i algorithm, default parameters, and the BLOSUM62 score matrix. The resulting MSA was manually inspected to identify a column in which the majority of residues were histidines, corresponding to the conserved phosphorylatable histidine. For sequences bearing a histidine at this position, the region spanning 5 amino acids upstream to 60 amino acids downstream of this residue was extracted, consistent with the structure of known HisKA domains in the Pfam database. As a final filtering step prior to profile generation, extracted sequences were aligned against known Pfam HisKA domains using HMMER and discarded if their bit score met or exceeded the gathering threshold (GA) of the corresponding profile.

### 2.3 Clustering, SEED and FULL profiles generation

These neighborhoods of conserved histidine residues were then used to generate SEED profiles. However, sequence diversity among these regions was too high to yield robust profiles. Sequences were therefore iteratively clustered, starting at 30% sequence identity and 80% coverage. When profiles derived from these clusters were too similar (see below), clustering was repeated at higher identity thresholds (40%, 50%, 55%, etc.). A cluster was retained for SEED profile construction if it contained ≥ 100 sequences, or ≥ 50 sequences if no cluster exceeded 100. Sequences within each cluster were then aligned using MAFFT with either the L-INS-i or FFT-NS-i algorithm, depending on cluster size.

Hidden Markov model (HMM) profiles were generated from the MSA of each cluster using HMMER. To identify potential similarities between profiles, each profile was aligned against all sequences grouped by SEED MSA. The minimum and maximum bit scores from these alignments were used to define the gathering (GA) and noise cutoff (NC) values. The GA of a profile was set to 20.0 by default and adjusted according to the following constraints. Let *s*_*i,jk*_ denote the bit score of the alignment of *S*_*jk*_, the *k*-th sequence from the *j*-th MSA (1 ≤ *k* ≤ |MSA_*j*_|), with the *i*-th profile and let GA_*i*_ denote the GA of that profile. In this case:

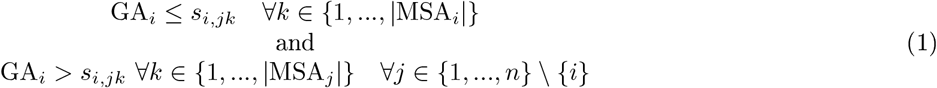

The NC of the *i*-th profile was set as GA_*i*_ − 0.1.

If a GA value could not be assigned to a profile under these constraints, the profile was deemed too general. If a set of profiles all overlapped with one another, then corresponding clusters were merged and a new profile was built. If some profiles lacked a GA, due to unmet requirements, and did not mutually overlap (e.g. A overlaps B, B overlaps C, but C does not overlap A), then their sequences were re-clustered at an increased identity threshold and new profiles were generated, after which the entire procedure was repeated to assign GA and NC values.

For each profile, a representative sequence was selected from the SEED alignment. Candidate representatives were required to have a valid UniProt ID and a total sequence length closest to the median length of all complete sequences in the SEED alignment. When two sequences were equally eligible, preference was given to the one with an entry in the STRING database (Szklarczyk et al., 2023). Throughout this article, each HisKA-like profile is referred to by the UniProt ID of its representative.

Once GA and NC values had been assigned to all profiles, FULL alignments were constructed. The new profiles were aligned against all iHKs that passed the filtering step (Figure 1, “Profiles and EC number filtering” box). Sequences were assigned to profiles based on their best-scoring match; when a sequence matched two profiles, it was assigned to the higher-scoring one, and the GA of the lower-scoring profile was adjusted to prevent future conflicts. FULL alignments were built using MAFFT with the L-INS-i or FFT-NS-i algorithm depending on the number of sequences, and FULL profiles were generated with HMMER. GA and NC values from the SEED profiles were transferred to the corresponding FULL profiles. All H-Boxes associated with iHKs that were not assigned to any of the new profiles were pooled and the workflow was resumed from the H-Box identification and conserved histidine steps, iterating until no histidine-conserved column could be identified in the H-Box alignment.

### 2.4 Profiles analysis

For cross-validation, we downloaded 566 proteins from SwissProt v2025_03 that matched the profile PS50109 but no HisKA Pfam profiles. We filtered out 423 sequences that matched known PFAM HisKA domains and, using the PyHMMER library (Larralde and Zeller, 2023), we aligned the resulting sequences against our profiles. Sequences yielding a match were then aligned against profiles SSF47384, SSF55890, SM00388, and PF14689 (HisKA-like profiles) for comparison. All profiles were downloaded from InterPro, except SM00388, for which no HMM file was available; instead, we retrieved the alignment from the SMART database and rebuilt the HMM profile using HMMER. The sequence of Lpl0330, an HK with an atypical HisKA domain not present in major databases, was downloaded from P2CS.

We performed a brief analysis of the genomic context of genes associated with our SEED profiles. The genomes of these genes were first annotated using eggNOG-mapper. For each gene in the SEED profiles, we then retrieved the COG categories and COG assignments of neighboring genes within their respective genomes, defined as consecutive genes separated by at most 400 nucleotides on either side. We computed a count matrix storing the occurrence of each COG category annotated in the genomic context of the 18 profiles, where rows correspond to profiles and columns to COG categories. From this count matrix, we derived an observed frequency matrix, which was subsequently visualized as a heatmap. Since a sequence can be assigned multiple COG categories, some observed frequencies exceeded 1. For instance, a sequence annotated with categories A and T was counted once under A and once under T. An analogous procedure was applied to individual COGs: we computed a count matrix of all COGs found in the genomic context, converted it to an observed frequency matrix, and generated a corresponding heatmap, retaining only COGs with an observed frequency *>* 0.05.

We used AlphaFold2-Multimer (Evans et al., 2021) version 2.3.1 to predict the homodimeric structure of each profile representative. The maximum template date was set to 2022-01-01. The database releases used by AlphaFold2 were as follows: UniRef90 (2022_04), MGnify (2022_05), PDB_mmCIF (February 14, 2023), PDB_seqres (January 13, 2023), Uniclust30 (2018_08), and UniProt (2022_04). For each predicted structure, we recorded the pLDDT, interface PAE, pTM, and ipTM scores.

For comparison with experimentally determined HisKA domain structures, we retrieved from InterPro (entry PF00512) the following reference structures: the homodimeric domain of the sensor HK EnvZ from *Escherichia coli* (PDB entry 1JOY), the DHp domain of PhoR (5UKV), the chimeric Af1503 HAMP—EnvZ DHp homodimer (2LFR), and the ERS1 dimerization and histidine phosphotransfer domain (4MT8). The region corresponding to the HisKA domain was extracted from the predicted structure of each profile representative. Pairwise structural alignments between these HisKA-like structures and the four reference HisKA structures were then performed using TM-align (Zhang, 2005). RMSD and TM-score were used as measures of structural similarity.

We also performed additional structural analyses. First, we searched the CATH, ECOD (Schaeffer et al., 2025), and TED (Lau et al., 2024) databases for structural classifications of the regions corresponding to our HisKA-like profiles. We then investigated whether the relative positions and orientations of the HisKA-like and HATPase domains were similar to those observed in experimentally determined HisKA—HATPase structures. For this purpose, we retrieved from the PDB 19 homodimeric structures containing HisKA and HATPase domains (4BIX, 5IDJ, 4Q20, 4U7O, 6E95, 9JFJ, 4JAU, 2C2A, 4BIU, 3D36, 4CB0, 6DK7, 6DK8, 4BIW, 4ZKI, 5C93, 4BIY, 6E52, and 4BIZ). These structures were aligned using TM-align against the HisKA-like—HATPase complexes previously predicted with AlphaFold2-Multimer. RMSD and TM-score were again used to quantify structural similarity.

We next examined the structural environment surrounding the predicted phosphorylatable histidine residue in our dimer models. Following the method proposed by Chen et al. (Chen et al., 2017), we compared the local structural environments (LSEs) of the predicted phosphorylatable histidine residues across the 18 representatives. Chen et al. introduced a scoring function to quantify the similarity between two LSEs. This score behaves like a distance measure, such that similar LSEs yield lower scores than dissimilar LSEs. As in the original study by Chen et al., we used a threshold of 0.5 to determine whether two LSEs were considered similar. Finally, for each HisKA-like domain, we compared the local structural environment of the histidine residue predicted to be phosphorylatable with those of the other histidine residues present in the same domain, when applicable.

To assess whether our profiles would spuriously align with non-HK proteins, we constructed a negative dataset of non-HK sequences. From InterPro, we downloaded all data listed under the “Domain Architectures” tab for five HATPase Pfam domains (PF02518, PF13581, PF13589, PF13749 and PF14501). Each domain architecture was characterized by a UniProt representative, a list of domains, and their positions within the representative sequence. Architectures containing a known HisKA or HisKA-like domain from the Pfam database were excluded. We then ran eggNOG-mapper on the representative sequences of the remaining architectures to obtain their EC numbers, and retained only sequences assigned an EC number indicative of enzymatic activity inconsistent with typical HKs. Sequences yielding matches with HisKA domain in SMART, SuperFamily, CDD, PROSITE, or CATH-Gene3D (Supplementary Table S1) were removed, and the resulting sequences were used to construct the negative dataset. Our profiles were then aligned against these sequences using PyHMMER.

We selected 41 well-studied organisms to evaluate the utility of our profiles in applied research. These organisms, along with the accession numbers of their reference genomes, are listed in Supplementary Table S7. Since the sequences used to build our SEED and FULL profiles derive exclusively from prokaryotic genomes, we restricted our selection to bacteria and archaea. Using PyHMMER, we aligned our profiles against the proteomes of these genomes and retrieved all matches above the GA threshold of each profile. We also assessed the proportion of sequences annotated as hypothetical protein among the resulting matches, based on the annotations provided in the GFF files of each genome.

## 3 Results

### 3.1 Eighteen probable HisKA HMM profiles

We started our analysis with 869 964 iHK sequences from bacterial, archaeal, fungal, and plant genomes, each carrying at least two domains: an HATPase domain and one additional domain (either C-terminal or N-terminal) . After the filtering and clustering steps described in section 2.1, we retained 92 334 sequences from bacterial genomes, 1 770 from archaeal genomes, 99 from fungal genomes, and 70 from plant genomes. We retrieved 32 785 matches with the PROSITE profile PS50109. Supplementary Figure S1 shows the distribution of match lengths against PS50109, which has a total length of 217 residues. With the exception of bacterial sequences, most sequences matched PS50109 across its full length, suggesting that they do harbor a HisKA-like domain that is at least partially detected by PS50109. For bacterial sequences, the failure to match PS50109 across its full length may indicate that their HisKA-like domains are too divergent from the one described by PS50109. We identified 14 591 H-Boxes and performed five rounds of clustering and alignment to identify novel HisKA-like domains.

The first round yielded nine profiles from clusters of at least 100 sequences, requiring five iterations of clustering to meet the criteria for defining GA and NC thresholds. Among these nine profiles, two were obtained by merging two clusters and one by merging three clusters, with clustering sequence identity reaching up to 55%. For subsequent rounds, the minimum cluster size was set to 50, as no cluster reached 100 sequences. The second round led to the identification of two additional profiles, requiring two iterations of clustering, the last at 50% identity. The third round produced two new profiles after clustering at 30% sequence identity. The fourth round yielded five new profiles after a single clustering step at 30% sequence identity. The fifth round was the last, as it was no longer possible to visually identify a column of conserved histidine residues in the H-Box MSA.

SEED and FULL HMM profiles were generated as described in section 2.3. By construction, the predicted phor-phorylatable histidine residue is always at position 6 in these profiles (since we build them using the —5/+60 neighborhoods of conserved histidine residues) . The FULL profiles collectively span 35 003 iHKs, with individual profiles matching between 82 and 25 876 sequences. All profiles were built exclusively from bacterial or archaeal sequences, as no plant or fungal sequences were included in any SEED or FULL profile. Supplementary Table S3 lists the 18 profiles along with the UniProt ID of their representative, the STRING ID where available, and a consensus sequence of the conserved histidine neighborhood. Sequence logos for the neighborhood of the conserved histidine residue in each profile are shown in Supplementary Figures S3 to S20.

### 3.2 Annotation transfer

We retrieved 566 protein sequences from SwissProt annotated with the PROSITE profile PS50109. Among these, 143 had no known Pfam HisKA domain annotated. Aligning these sequences against our profiles yielded 27 matches across three profiles: F4GBN6, A0A221P3F7 and A0A7X9X266, this with 12, 11 and 4 hits, respectively. For all matching sequences, the position of the phosphorylatable histidine residue had been manually curated by SwissProt annotators. Our profiles correctly identified the phosphorylatable residue in every case (Supplementary Table S4).

These sequences also matched two SuperFamily profiles: SSF47384 and SSF55890, which characterize the homodimeric domain of signal transduction histidine kinases and the sporulation response regulatory protein Spo0B, respectively. Some additionally matched the Pfam entry PF14689, associated with Spo0B, and the SMART entry SM00388 (the HisKA domain in SMART) . Since all matching sequences possess a conserved histidine residue, we compared the alignments obtained with these established profiles against those obtained with our HisKA-like profiles.

For the 12 sequences matching both F4GBN6 and SSF47384, alignment scores with F4GBN6 were consistently higher than with SSF47384, as shown in Supplementary Table S5. Supplementary Figure S2 shows the alignment of the H-Boxes from these 12 sequences alongside the sequence logos of profiles SSF47384 and F4GBN6. Results were more nuanced for the 11 sequences matching PF14689, SSF55890, and profile A0A221P3F7. As shown in Supplementary Table S5, half of the sequences scored higher with A0A221P3F7 and the other half with SSF55890, while PF14689 consistently yielded the lowest scores. Finally, the four sequences that aligned with SSF47384, SM00388, and profile A0A7X9X266 all scored higher with A0A7X9X266 than with either of the other two profiles.

We also searched the literature for HKs harboring atypical HisKA domains. We identified Lpl0330 (Levet-Paulo et al., 2011), an HK whose HisKA domain was uncharacterized at the time of publication and remained absent from domain databases at the time of submission of this article. Aligning the sequence of Lpl0330 against our 18 profiles yielded one hit with profile A0A0H2MHX8. However, this hit identified a different histidine residue than the one experimentally characterized by Levet-Paulo et al. (His-296 vs. His-210, respectively). To investigate this discrepancy, we aligned all complete sequences forming profile A0A0H2MHX8 and found that two conserved histidine residues were present upstream of the HATPase domain. Our method had selected the more C-terminal histidine residue, whereas the true phosphorylatable residue appears to be the one located on the N-terminal side. Profile A0A0H2MHX8 was accordingly corrected by re-applying our methodology using this alternative histidine residue as the reference.

### 3.3 3D structures of the HisKA-like domains

As protein function is closely linked to protein structure, we compared the 3D structures of our profile representatives with those of experimentally determined HisKA domains. The HisKA domain is well characterized and consists of two α-helices, *α*1 and *α*2 (Marina et al., 2005), with the phosphorylatable histidine residue located approximately midway along the solvent-exposed surface of helix *α*1, as described by Marina et al. Due to computational resource constraints, we restricted this analysis to the profile representatives and used AlphaFold2-Multimer to predict their homodimeric 3D structures.

Figure 2 shows the pLDDT values along the first chain of each predicted dimer, with red rectangles indicating the regions corresponding to our HisKA-like profiles. The pLDDT values for the second chain were identical to those of the first chain. Table 1 lists the pTM and ipTM scores associated with each prediction, together with the pLDDT score of the predicted phosphorylatable histidine residue. The interface PAE for each predicted dimer is shown in Figures 3–5, with red boxes indicating the regions corresponding to the HisKA-like domains.

**Table 1:** pTM and ipTM values for each dimer, as well as the pLDDT score for the conserved histidine residue.

| Representative's ID | pTM | ipTM | Conserved histidine's pLDDT |
| --- | --- | --- | --- |
| A0A0H2MHX8 | 0.497 | 0.479 | 64.089 |
| A0A1G9BVG1 | 0.598 | 0.585 | 76.845 |
| A0A1I3H793 | 0.585 | 0.582 | 60.951 |
| A0A418Q055 | 0.524 | 0.503 | 58.269 |
| A0A6L9EDD3 | 0.506 | 0.495 | 66.345 |
| A0A8D6V7Y3 | 0.658 | 0.658 | 68.191 |
| B3E1V6 | 0.846 | 0.817 | 74.176 |
| C6BWG9 | 0.471 | 0.462 | 60.742 |
| J1L467 | 0.532 | 0.526 | 74.081 |
| R0D498 | 0.684 | 0.670 | 69.729 |
| A0A1G7WDI6 | 0.467 | 0.450 | 62.194 |
| A0A1H9IBY7 | 0.451 | 0.435 | 91.185 |
| A0A221P3F7 | 0.515 | 0.494 | 60.436 |
| A0A6H0KTZ1 | 0.715 | 0.709 | 91.905 |
| A0A7X9X266 | 0.567 | 0.562 | 59.974 |
| A0A936YZK5 | 0.548 | 0.538 | 62.310 |
| B9LU29 | 0.707 | 0.717 | 82.506 |
| F4GBN6 | 0.592 | 0.569 | 65.050 |

**Figure 2:**
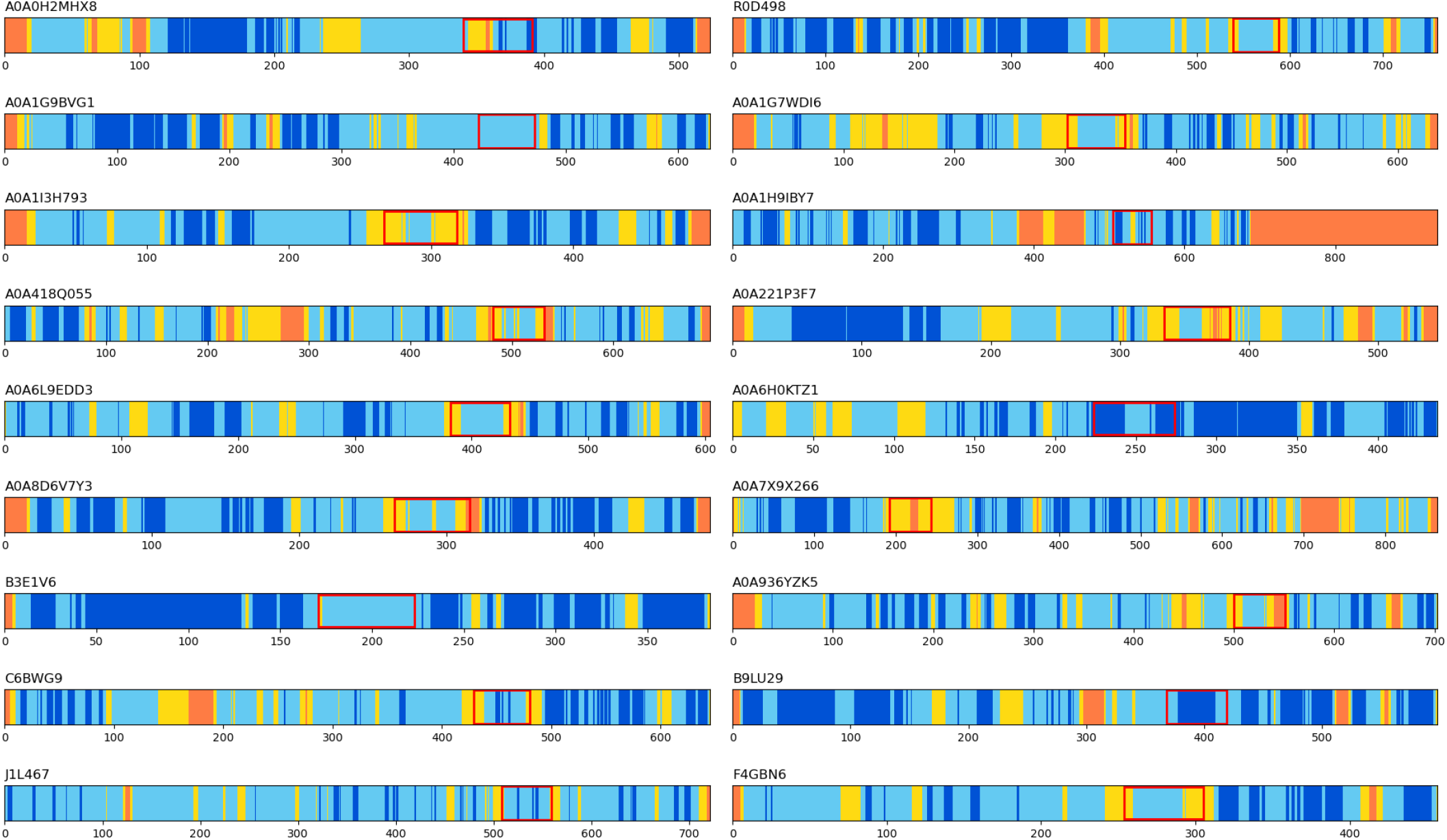
pLDDT values for the first chain of each dimer. The values for the second chains are identical to those displayed here. Red rectangles correspond to the HisKA-like domains.

**Figure 3:**
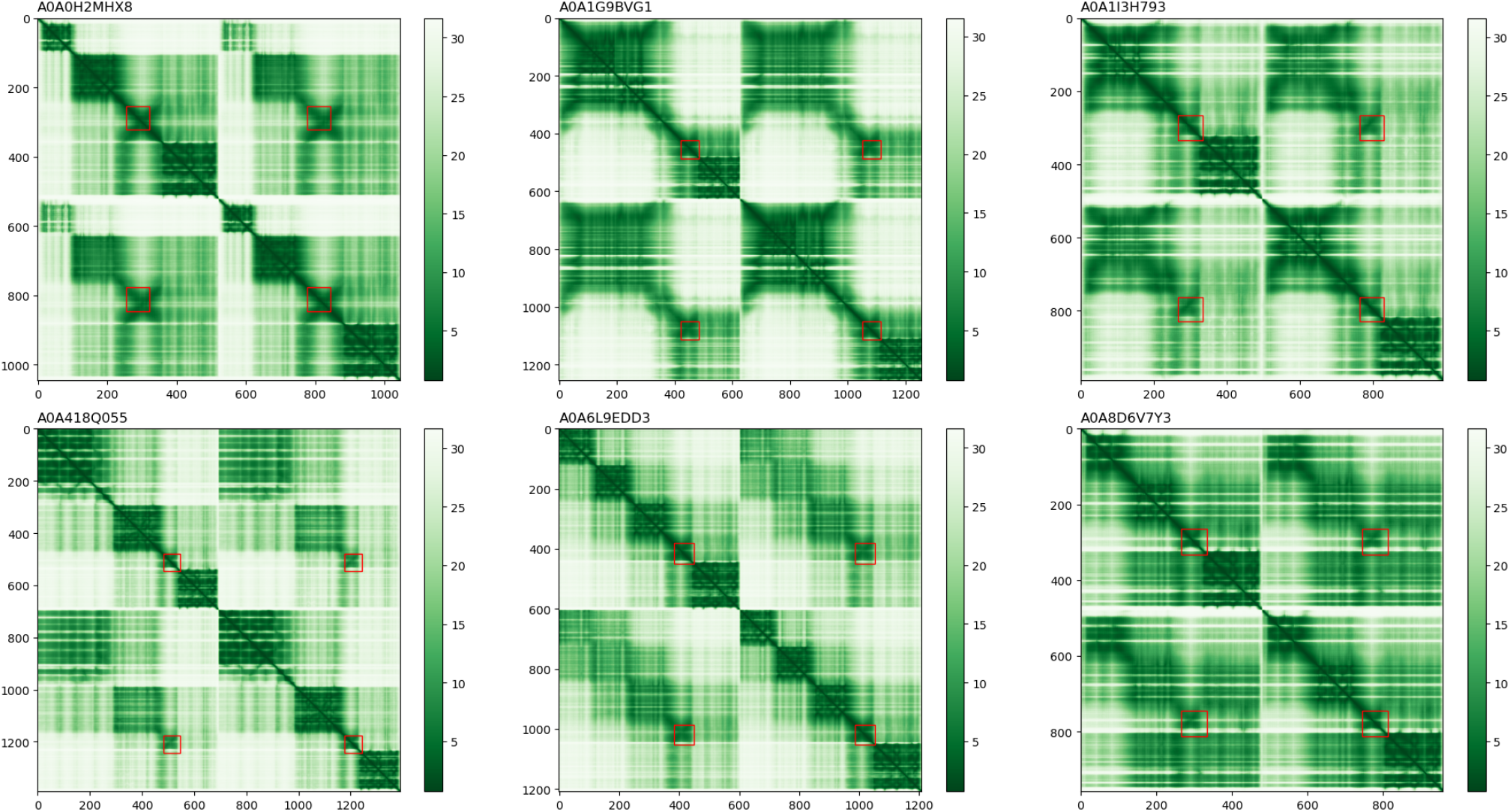
Interface PAE for six of the eighteen dimer models. Red rectangles highlight the regions aligning with our HisKA-like profiles.

**Figure 4:**
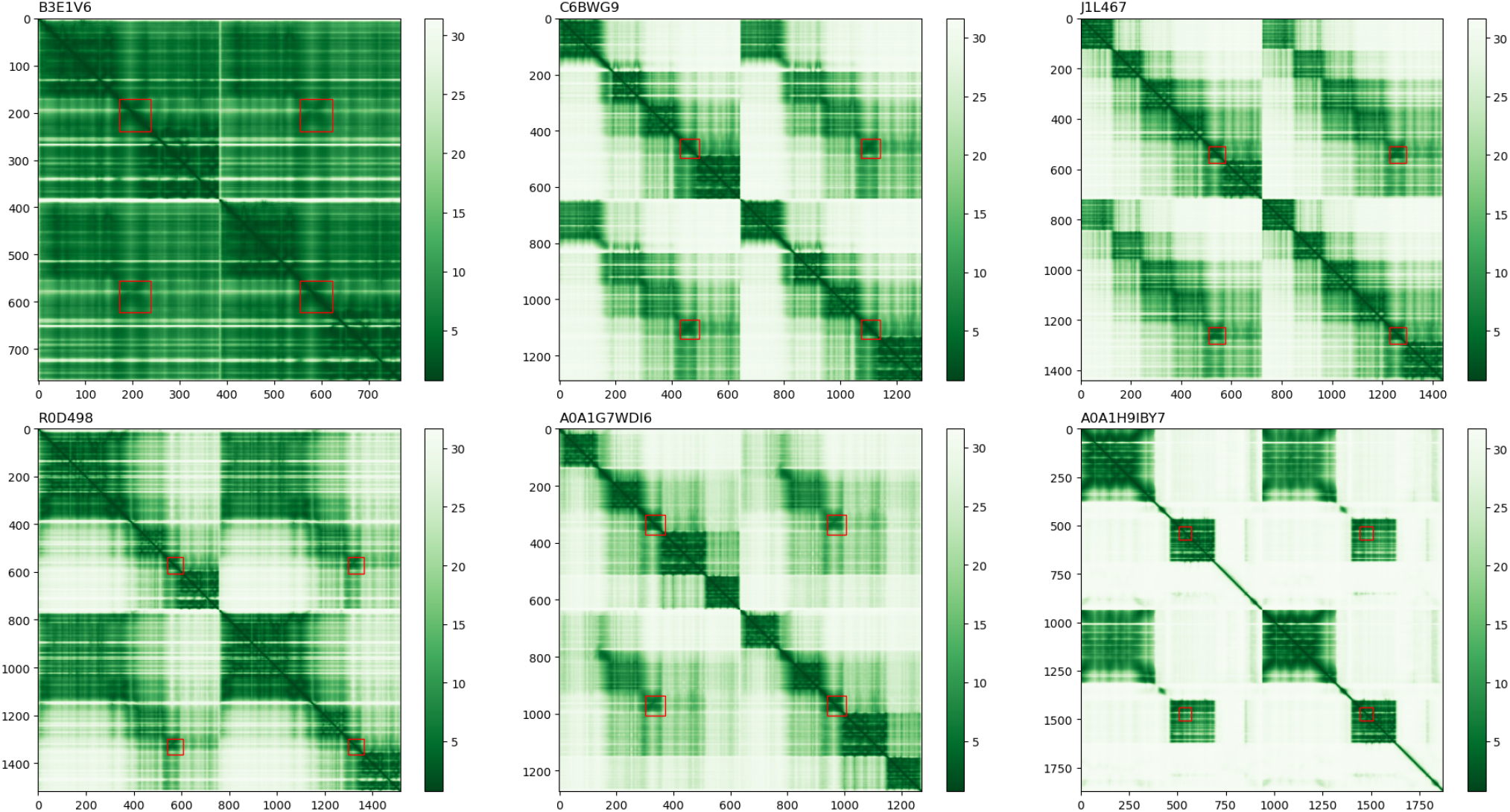
Interface PAE for six of the eighteen dimer models. Red rectangles highlight the regions aligning with our HisKA-like profiles.

**Figure 5:**
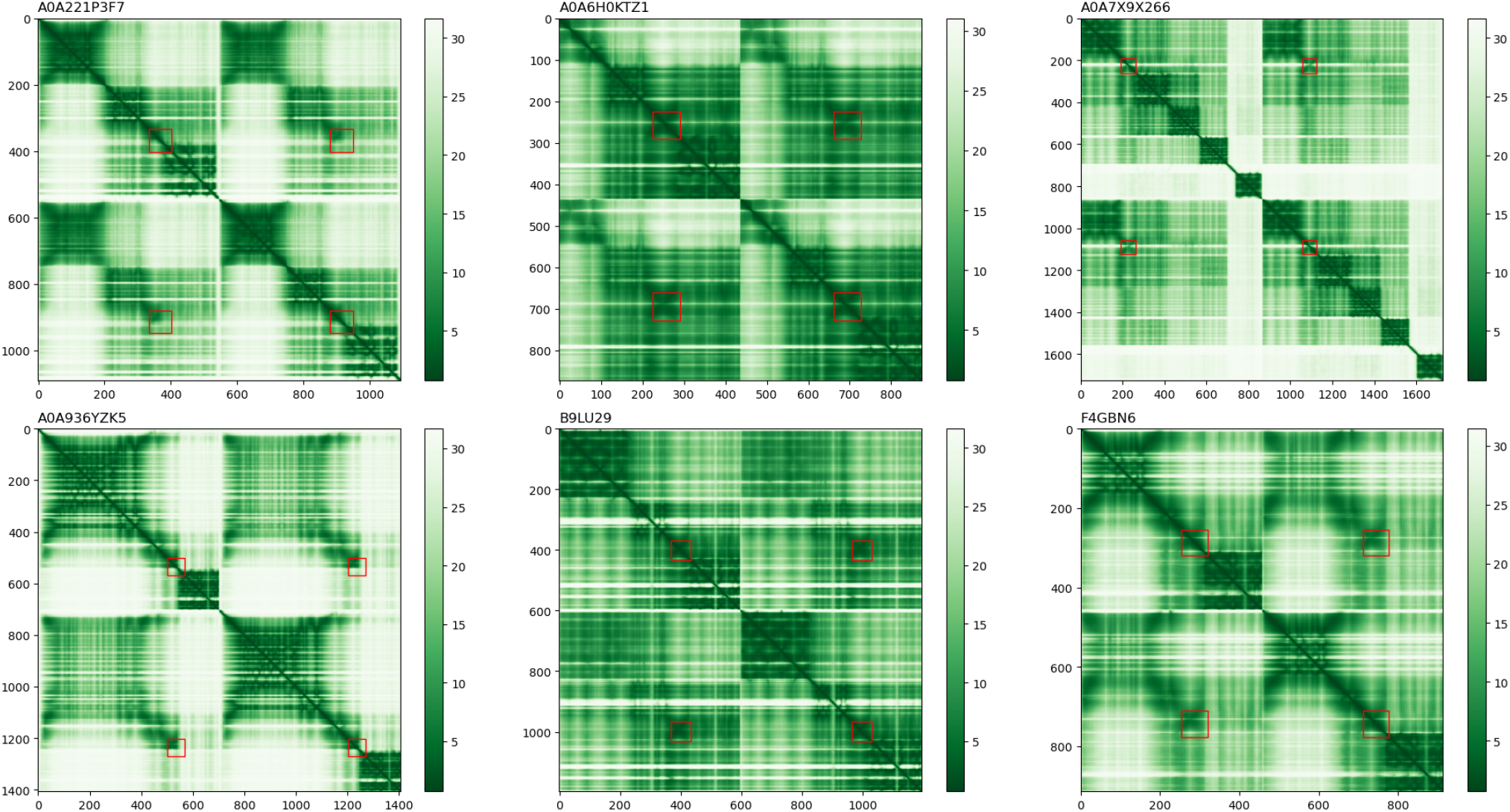
Interface PAE for six of the eighteen dimer models. Red rectangles highlight the regions aligning with our HisKA-like profiles.

Figure 6 shows the predicted homodimeric structure of the representative of profile A0A6H0KTZ1 and compares it with the experimentally determined structure of the EnvZ homodimeric domain (Tomomori et al., 1999).

**Figure 6:**
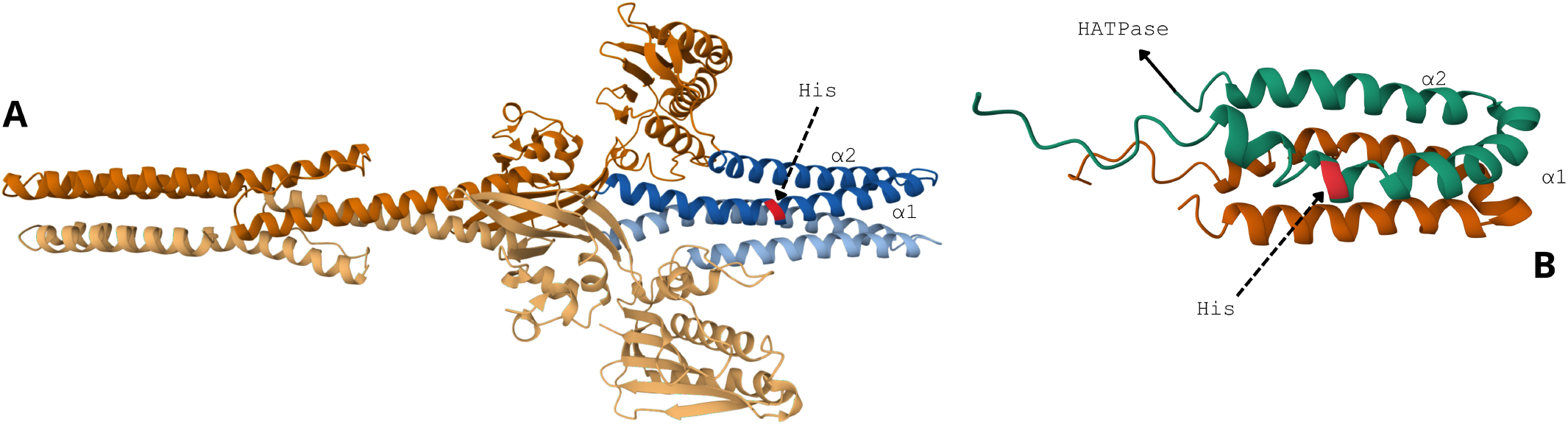
**A**: Predicted homodimeric 3D structure of the representative sequence of profile A0A6H0KTZ1, generated using AlphaFold2-Multimer. The two chains of the homodimer are distinguished by color shade. The two α-helices of the HisKA-like domain are highlighted in blue. The conserved histidine residue of the first chain is highlighted in red. **B**: Structure of the homodimeric domain of EnvZ (*Escherichia coli*), downloaded from InterPro (originally determined by multidimensional NMR). Using the green chain as spatial reference, the first *α*-helix is at the bottom, with the histidine residue of interest highlighted in red.

Structures of all other profile representatives are available in the data archive link provided in section 6.3. As shown in Figure 6, the structure of the HisKA-like domain is visually similar to that of the EnvZ homodimeric domain. The histidine residue predicted to be phosphorylatable by our profile is correctly positioned within the first *α*-helix. Similar results were observed for other profile representatives, with the exception of profile A0A1H9IBY7, where this histidine is positioned in the second *α*-helix.

Regarding the output metrics produced by AlphaFold2-Multimer, and starting with the pLDDT, we can see on Figure 2 that there is a marked contrast among the HisKA-like regions in the dimers. For example, B9LU29 display good overall pLDDT scores and others, in particular A0A7X9X266, are displaying lower values. In many cases, the extremities of the HisKA-like domains have pLDDT values below 70, thus indicating low-confidence regions. Regarding the pLDDT of the conserved histidine residue, Table 1 show that in most cases (12 out of 18) it is below 70. In addition, the pTM and ipTM scores in the same Table do not indicates high confidence for most models. In particular ipTM scores, which should be above 0.8 ideally, are generally around 0.5. With pTM scores also around 0.5, the overall structure of each dimer is predicted with moderate confidence, whereas the ipTM scores indicate that the relationships between the different subunits are not well defined. However three dimers stand out – B3E1V6, A0A6H0KTZ1, and B9LU29– with pTM and ipTM scores above 0.7. The histidine residue in their respective HisKA-like domains have the highest pLDDT scores among all others.

Concerning the interface PAE, most plots (except for B3E1V6, A0A6H0KTZ1, and B9LU29) reflect the same pattern as the ipTM scores, namely low-confidence relationships between the subunits of each dimer. Nevertheless, in each model, our HisKA-like profiles align with regions that appear to form well-defined folds. By looking at the PAEs and the red rectangles corresponding to the HisKA-like domains, it appears that they overlap slightly the N-terminal extremities of the HATPase domains. In this regard, we re-generated new HisKA-like profiles, this time by taking 45 aa after the conserved histidine (instead of 60). These new versions of the profiles are given alongside their first version. Supplementary Figures S21 to S23 display the interface PAE of each dimer with the second version of the profiles mapped using red rectangles.

Overall, the metrics produced by AlphaFold2-Multimer indicate that these models should be taken with caution. This is mainly due to the fact that these models are incomplete, as an HK most of the time dimerize upon signal recognition (e.g ligand binding) and this signal is not known here. However, the PAE plots show that the regions aligning with our HisKA-like profiles are well defined in the dimers (green blocks on and off the diagonal). We also looked at the monomeric version of these structures (available on UniProt) and these regions have higher pLDDT scores, with similar folds typical of HisKA domains. Supplementary Table S6 displays these pLDDT scores. With this in mind, we decided to continue the structural analysis on the predicted structure of our HisKA-like domains.

We extended the structural analysis to additional HisKA reference structures using a more quantitative approach. For each representative, we extracted the part of the structure corresponding to the HisKA-like domain. Using TM-Align, we performed pairwise structural alignments between the structures of these HisKA-like domains and four experimentally validated HisKA reference structures from the PDB database: 2LFR (chimeric Af1503 HAMP –EnvZ DHp homodimer), 4MT8 (ERS1 dimerization and histidine phosphotransfer domain from *Arabidopsis thaliana*), 1JOY (EnvZ homodimeric domain from *E. coli*), and 5UKV (DHp domain of PhoR). Figure 7 shows the distributions of RMSD and TM-scores resulting from these pairwise alignments for each reference structure.

**Figure 7:**
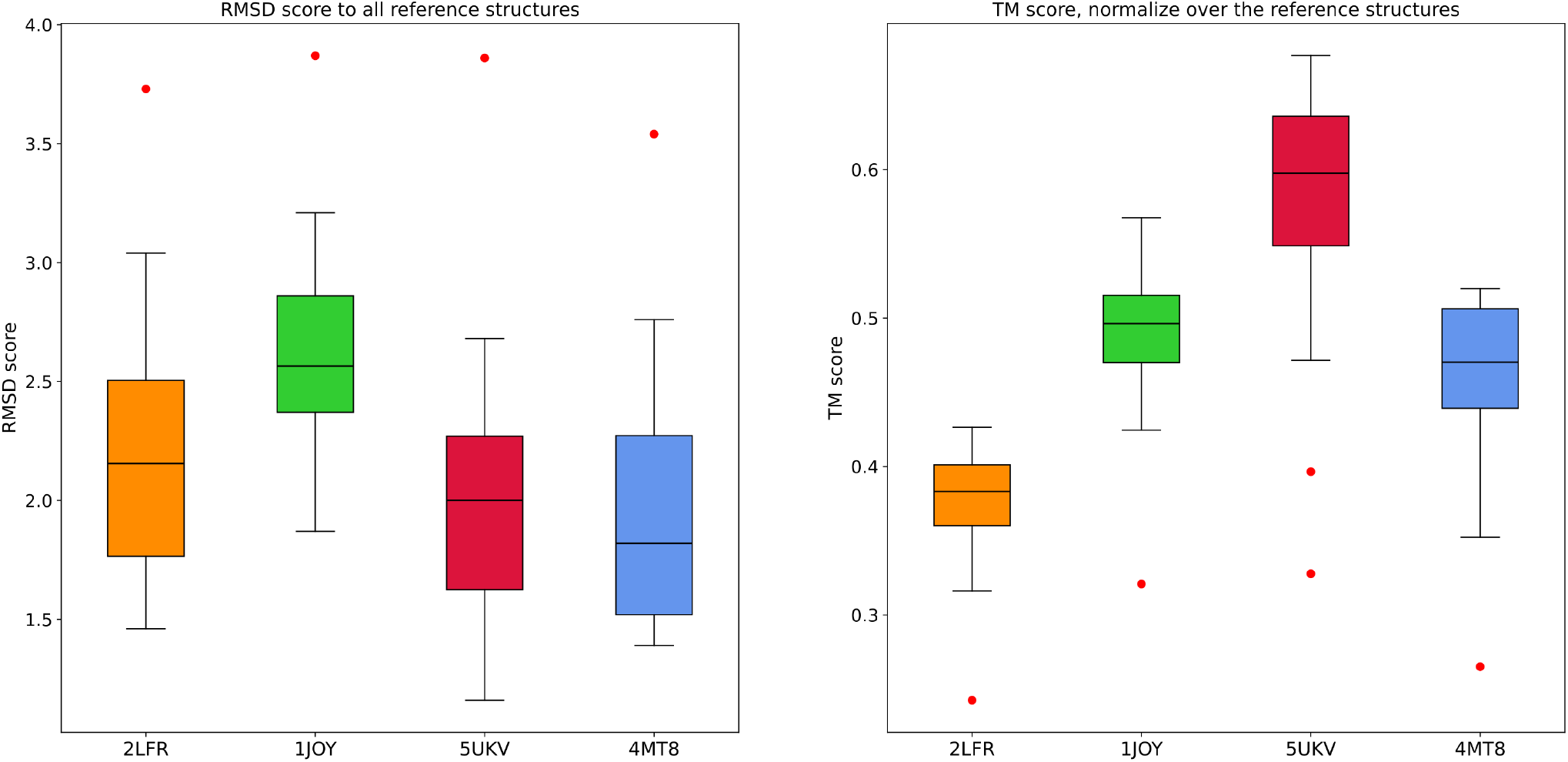
Distribution of RMSD and TM-scores resulting from the pairwise alignment between each reference structure against all HisKA-like representative structures. From left to right, the PDB IDs on the x-axis correspond to: EnvZ homodimeric domain, DHp domain of PhoR, chimeric Af1503 HAMP–EnvZ DHp homodimer and ERS1 dimerization and histidine phosphotransfer domain from Arabidopsis thaliana.

RMSD values were approximately 2.0 across all reference structures, with the exception of 1JOY for which they were slightly higher at around 2.5. TM-scores were distributed around 0.5 for the EnvZ homodimeric domain (1JOY) and the ERS1 dimerization and histidine phosphotransfer domain (4MT8). Alignments against the DHp domain of PhoR (5UKV) yielded higher TM-scores, centered around 0.6. The lowest TM-scores were observed for 2LFR, most likely because this structure encompasses both a HisKA and a HAMP domain, making it considerably larger. Profile A0A1H9IBY7 stood out as an outlier in each boxplot. A second outlier, for the TM-score against 5UKV, was profile A0A936YZK5. Overall, the closest reference structure appears to be 5UKV, with both RMSD and TM-score values indicative of a good degree of structural similarity.

Using databases ECOD, CATH and TED we conducted structural classifications of our HisKA-like profiles. With the ECOD database, we used two different search options, the first based on BLAST (Camacho et al., 2009) and using the sequences of the representatives, the second based on Foldseek (Van Kempen et al., 2024) and using the predicted homodimeric structures of the representatives. We will only report matches concerning the HisKA-like profiles, as the presence of other domains is known (section 2.1).

The ECOD sequence search (on ecod100_af2_pdb with E-value cut-off at 0.01) did not reports many matches beside ECOD family 605.1.1.1 (HisKA family) for A0A1G9BVG1 and B9LU29. We found matches with ECOD topology 605.1.1.0 (in which is found the HisKA family) for A0A1G9BVG1, A0A1H9IBY7, B9LU29, J1L467, R0D498. The lack of matches with the sequence search is due to the fact that ECOD families are build upon Pfam and that the iHKs we analyzed do not match any HisKA profiles in Pfam. The ECOD structure search (against ECOD representative domains, E-value cut-off at 0.01) yielded more matches, with family 605.1.1.1 found in A0A0H2MHX8, A0A1G7WDI6, A0A1G9BVG1, A0A6L9EDD3, B9LU29, C6BWG9, J1L467 and R0D498. Topology 605.1.1.0 was reported in all queries.

For the CATH classification, we performed a sequence-based search that did not produced many matches as well. We had two matches with the “Signal transduction histidine kinase, dimerisation/phosphotransfer (DHp)” domain (1.10.287.130) on A0A221P3F7 and F4GBN6 (respective FunFams entries FF11 and FF125 & FF123). We performed an additional structural classification using the TED database. In the TED database, predicted protein structures from UniProt are parsed to identify structural domains. Because each HisKA-like representative as a valid UniProt ID, we were able to retrieve their TED annotations (except for A0A8D6V7Y3 and A0A936YZK5 who were not in the TED database). Table 2 shows that, for 10 out of 18 profile representatives, the TED pipeline identified domains that coincide in position with our HisKA-like profiles. Moreover, for eight of them, a link to CATH topology 1.10.287 was established by the TED pipeline. This topology corresponds to the one containing CATH homology 1.10.287.130 (DHp domain). The A0A1G7WDI6 HisKA-like domain is associated with CATH topology 1.20.5, which corresponds to single *α*-helices but this may be an annotation error because the TED domain clearly spans two parallel *α*-helices.

**Table 2:** TED annotations that coincide with regions aligning with our HisKA-like profiles. This concerns 10 of the 18 profie representatives. CATH classification is reported when available.

| Representative's ID | TED domain start-end | HisKA-like profile start-end | CATH classification |
| --- | --- | --- | --- |
| A0A0H2MHX8 | 232-357 | 254-321 | - |
| A0A1G7WDI6 | 287-359 | 302-370 | 1.20.5 |
| A0A1G9BVG1 | 394-481 | 422-484 | 1.10.287 |
| A0A1I3H793 | 246-322 | 267-333 | 1.10.287 |
| A0A418Q055 | 486-533 | 481-546 | 1.10.287 |
| A0A6H0KTZ1 | 212-284 | 224-290 | 1.10.287 |
| A0A6L9EDD3 | 361-441 | 382-448 | 1.10.287 |
| C6BWG9 | 417-488 | 429-495 | 1.10.287 |
| F4GBN6 | 231-307 | 254-319 | 1.10.287 |
| J1L467 | 492-563 | 508-574 | 1.10.287 |

Using the method introduced by Chen et al. (Chen et al., 2017), we compared the LSE of the phosphorylatable histidine residues found in the 18 profile representatives. Chen et al. proposed a scoring function to quantify similarity between LSEs, with similar LSEs receiving a score close to 0. In our study, as in the initial study of Chen et al., we considered that two LSEs were similar if their score was below 0.5. Figure 8 displays the pairwise comparison of all the phosphorylatable histidine residues. We can see that the similarity scores are heterogeneous. For four representatives —J1L467, A0A1G7WDI6, F4GBN6 and A0A6H0KTZ1— the LSE of the target histidine residue is similar to the LSE of the target histidine in all the others representatives (similarity score always below 0.5). But regarding the rest of the representatives, we notice dissimilarities between target histidine. In particular, the representatives at the top of the heatmap have a phosphorylatable histidine residue whose LSE differs from those of the others (A0A1H9IBY7 is a special case because others analysis put its HisKA-like characteristics in doubt) . However for each representative there is always at least one match with a similarity score below 0.5.

**Figure 8:**
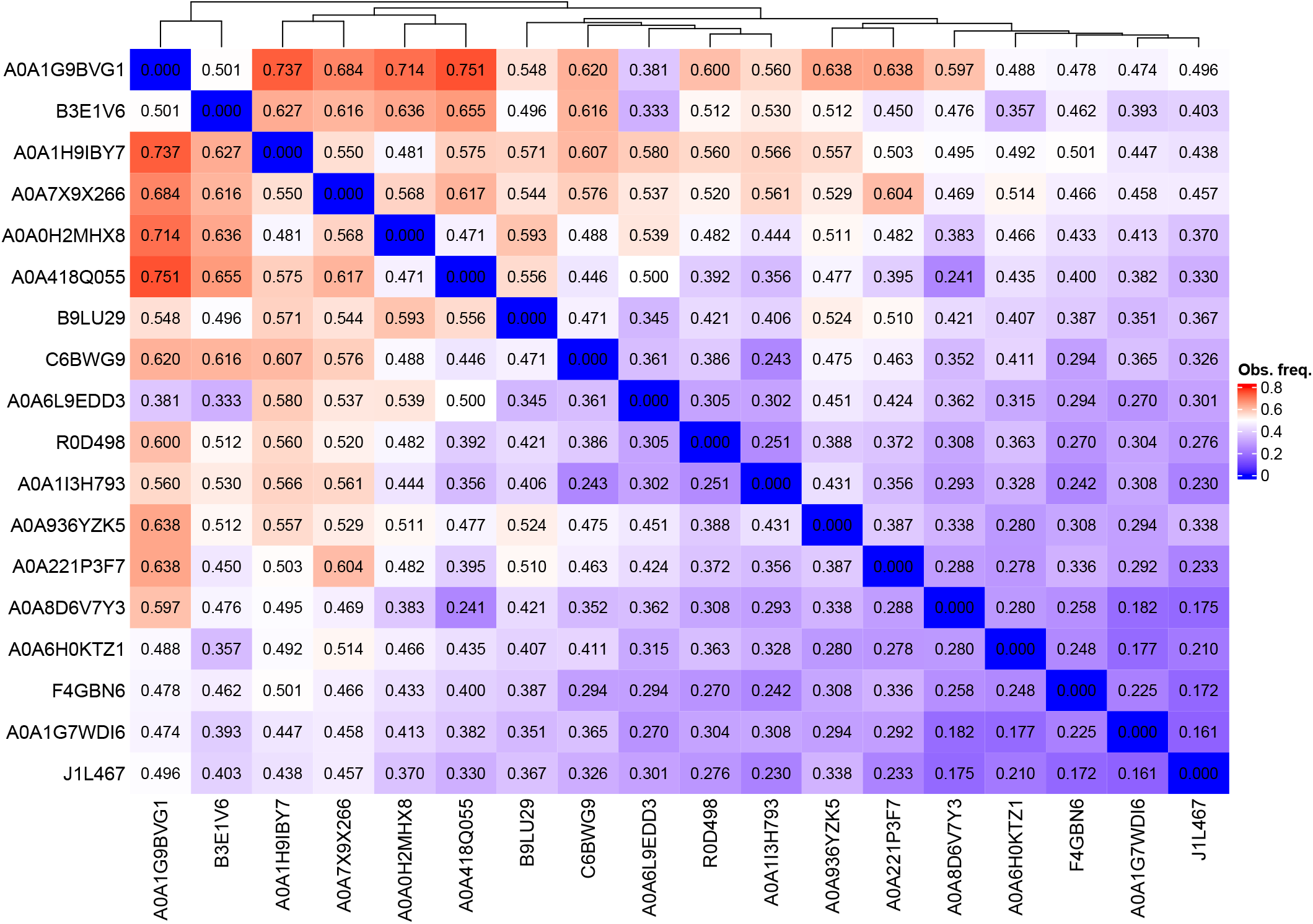
Pairwise comparison of the LSE of each conserved histidine residue from the 18 HisKA-like profiles. Dendogram at the top represent a hierarchical clustering of the profiles.

We also looked at the structural features of the predicted phosphorylatable histidine residues to see if they could distinguish them from other histidine residues within the same HisKA-like domains. The main structural feature of the phosphorylatable histidine residue is its position within the two parallel *α*-helices that form the HisKA domain. As explained previously, the phosphorylatable histidine residue is supposed to be in the middle of the first *α*-helix. We can also look at the solvent-accessible surface areas (SASA) of the histidine residues, as the phosphorylatable histidine will have a high SASA value. Figure 9 displays the SASA values of histidine residues found in the HisKA-like regions of representatives, for those containing more than one histidine residue in these regions.

**Figure 9:**
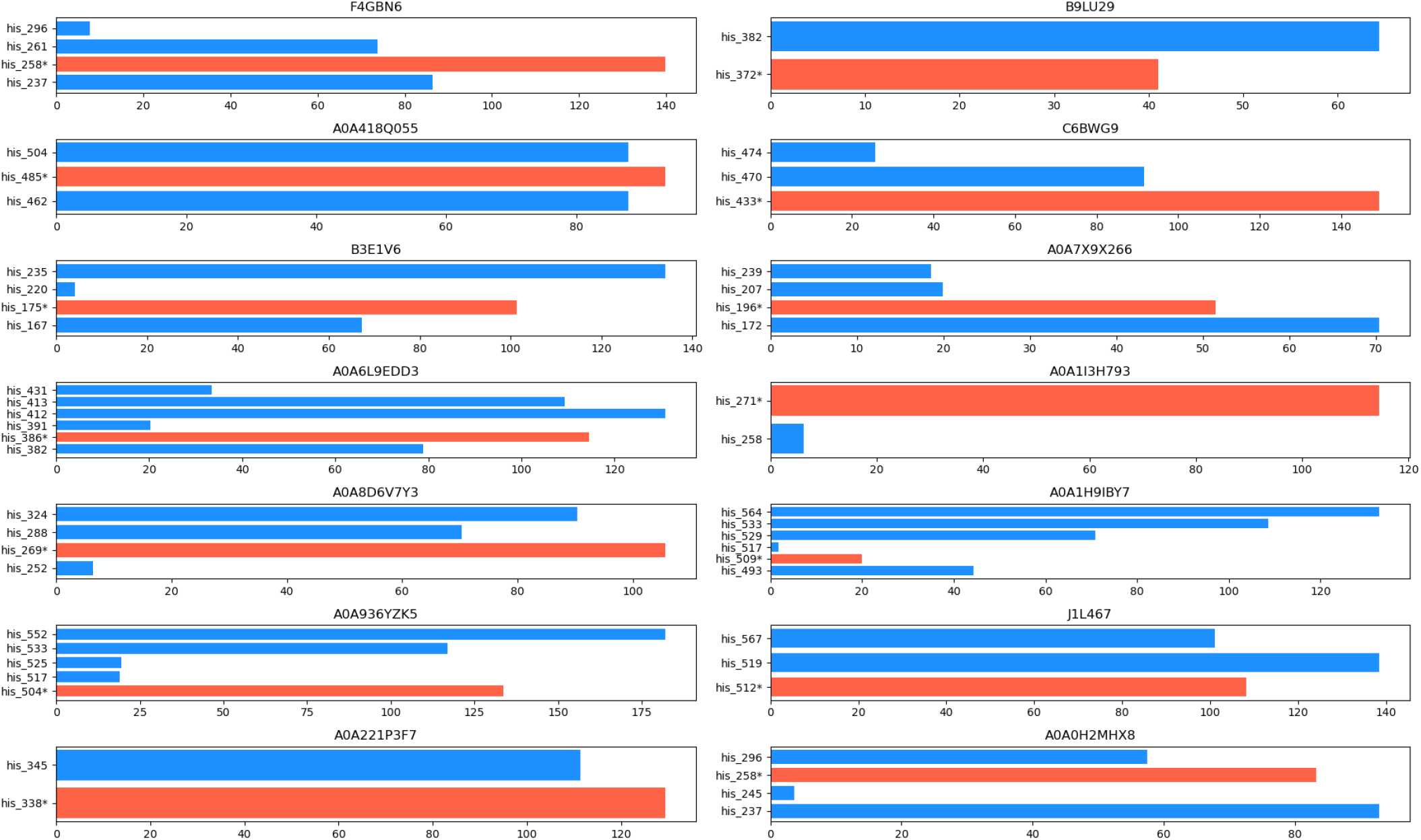
SASA values of the histidine residues in the HisKA-like regions. We only display cases in which there is more than one histidine residue in the HisKA-like region. Red bars correspond to the histidine residue that is predicted as phosphorylatable by our profiles.

For half of the representatives, the predicted phosphorylatable histidine residue has a higher SASA value than the other histidine residues. For the other half —B3E1V6, A0A6L9EDD3, A0A936YZK5, B9LU29, A0A1H9IBY7, J1L467 and A0A0H2MHX8— the predicted phosphorylatable histidine residue ranks second in terms of SASA value. One exception is A0A1H9IBY7, for which the target histidine has among the lowest SASA value. A0A1H9IBY7 has already been flagged as questionable, as the conserved histidine residue of its associated HisKA-like domain is located in the second *α*-helix. For other representatives, the histidine residues with higher SASA value than the predicted phosphorylatable histidine residues are never found in the middle of the first *α*-helix, sometimes even found in the second *α*-helix. In conclusion, the conserved histidine residues of our profiles are always the only histidine residues located in the middle of the first *α*-helix with the highest SASA value (except for A0A1H9IBY7) .

Lastly, we compared the HisKA_like-HATPase complexes identified in the predicted structures of our representatives with the HisKA-HATPase complexes observed in the experimental structures of HKs. We used TM-Align to align our HisKA_like-HATPase dimer against 19 HisKA-HATPase dimer from the PDB. Figure 10 shows the resulting

**Figure 10:**
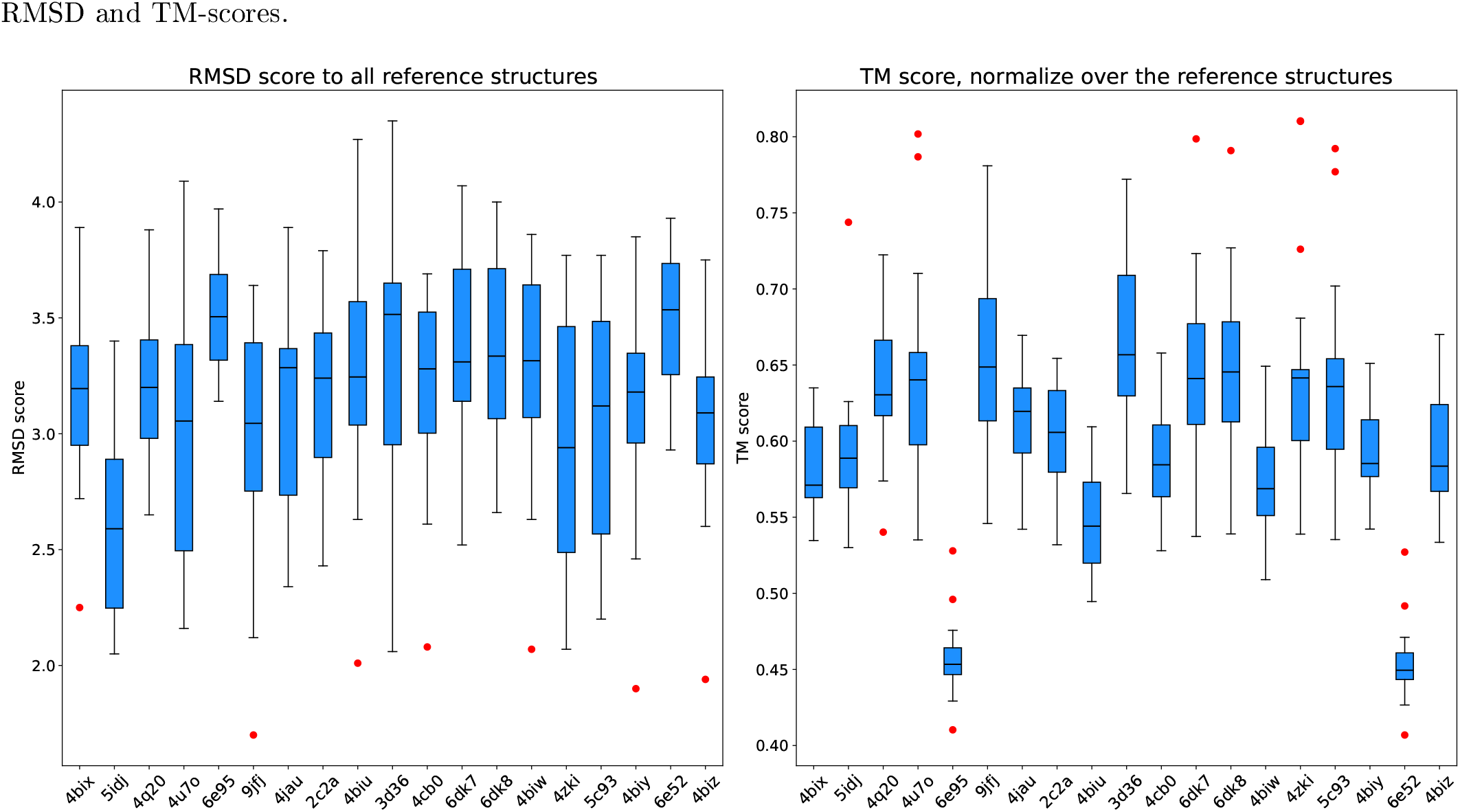
Distribution of RMSD values and TM-scores resulting from structural alignments between PDB reference HisKA-HATPase complexes and the HisKA_like-HATPase complexes of our representatives. The x-axis corresponds to the PDB ID of the reference structure.

Resulting RMSD are higher than the one obtained with previous HisKA-like/HisKA structural alignments but are still acceptable. This is mainly due to the larger size of the aligned structures, leading to more residues being misaligned. However the TM-scores are higher on average than the ones displayed in Figure 7, and are around 0.6 or above for most HisKA-HATPase references. We can see that two reference structures yield poor results for both RMSD and TM-score : 6e95 and 6e52. These two structures correspond to a leucine zipper domain fused to a HisKA-HATPase complex. The presence of that additional domain explain the low alignment scores. Overall, it appears that the relationship between the HisKA-like and HATPase domains in our predicted structures is similar to that observed between the HisKA and HATPase domains in experimental structures.

### 3.4 Gene context implies regulation activity

One of our analyses focused on the genomic context of genes present in our 18 SEED alignments. We used eggNOG-mapper to annotate the corresponding genomes and retrieved COG assignments and COG categories for neighboring genes. The eggNOG database was chosen for several reasons: it provides a robust framework for functional annotation, is built with taxonomic diversity in mind, and is regularly updated. Our SEED alignments contained 9 471 genes in total. For each gene, we determined its genomic context by identifying neighboring genes within the same genome, yielding more than 30 000 genes co-occurring in the same genomic neighborhoods. Of these, 2 157 lacked any COG annotation (neither COG assignment nor category) and 759 had a COG category but no COG assignment.

We first examined COG categories, as they provide a broad summary of the functional activities represented in these genomic neighborhoods. As described in section 2.4, we computed a matrix of observed frequencies of COG categories for the genomic context of each profile and generated the heatmap displayed in Figure 11. The most frequently observed categories were T, S, and K, corresponding to “Signal transduction mechanisms”, “Function unknown”, and “Transcription”, respectively. Profiles lacking category T in their genomic context tended to show a higher representation of category K instead, with the exception of profile B9LU29. The least represented categories were A, B, W, and Z (“RNA processing and modification”, “Chromatin structure and dynamics”, “Extracellular structures”, and “Cytoskeleton”, respectively), with no profile reaching an observed frequency ≥ 0.1. Category A was particularly scarce in the genomic context of profile A0A418Q055.

**Figure 11:**
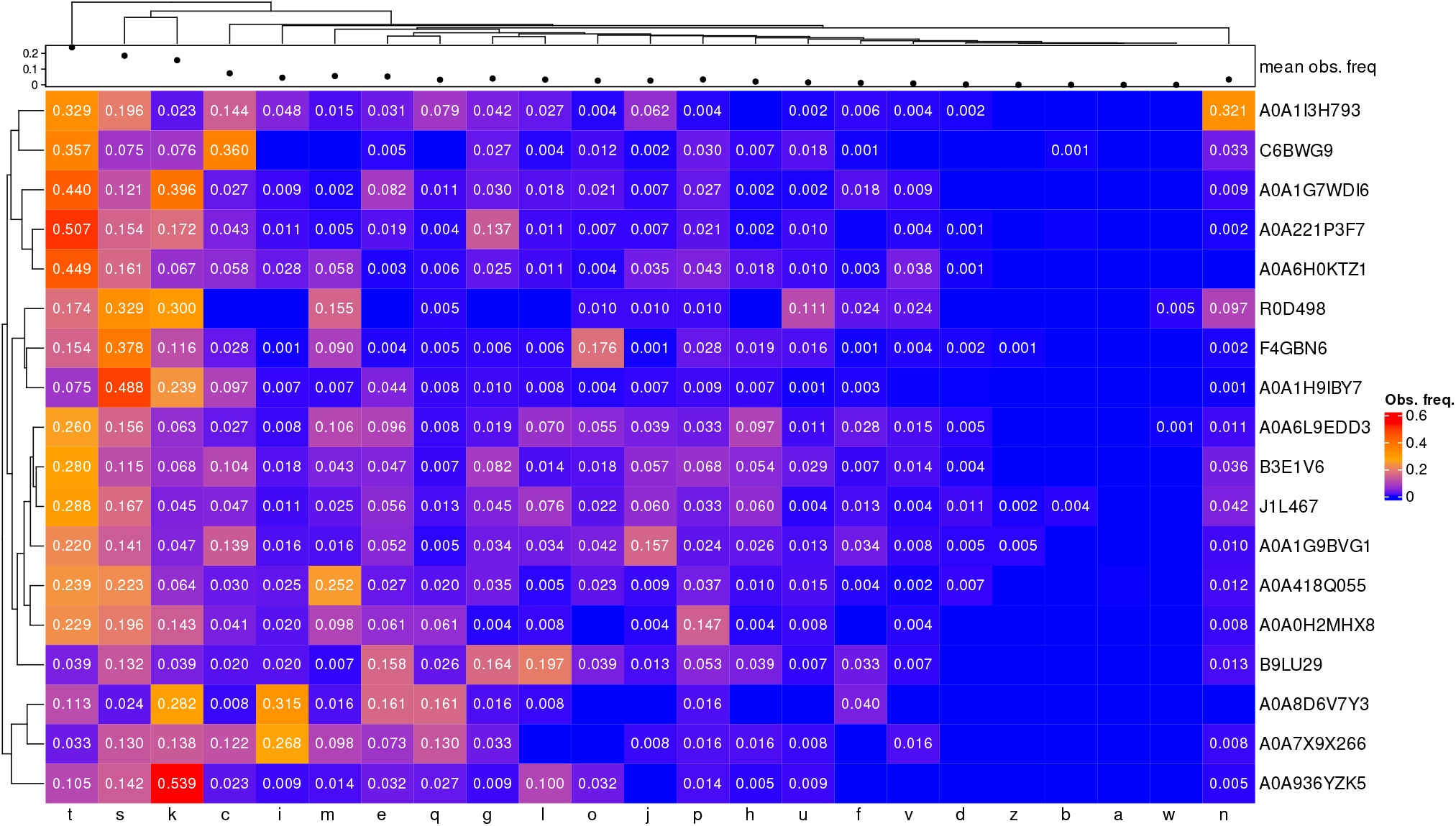
Heatmap of the observed frequency of COG categories in the genomic context of all 18 profiles. The bar plot at the top displays the mean observed frequency of each COG category across all profiles. The dendrogram on the left shows similarities between profiles, and the dendrogram at the top shows similarities between COG categories. Values below 0. 001 are not reported.

Visual inspection of the heatmap in Figure 11 suggests that some profiles share similarities in their COG category composition. To assess whether the profiles also share more specific regulatory functions, we generated a second heatmap focusing on individual COG assignments, displayed in Figure 12. Given the large number of annotated COGs (1 636), we retained only those with an observed frequency *>* 0.05 in at least one genomic context, reducing the set to 52 COGs.

**Figure 12:**
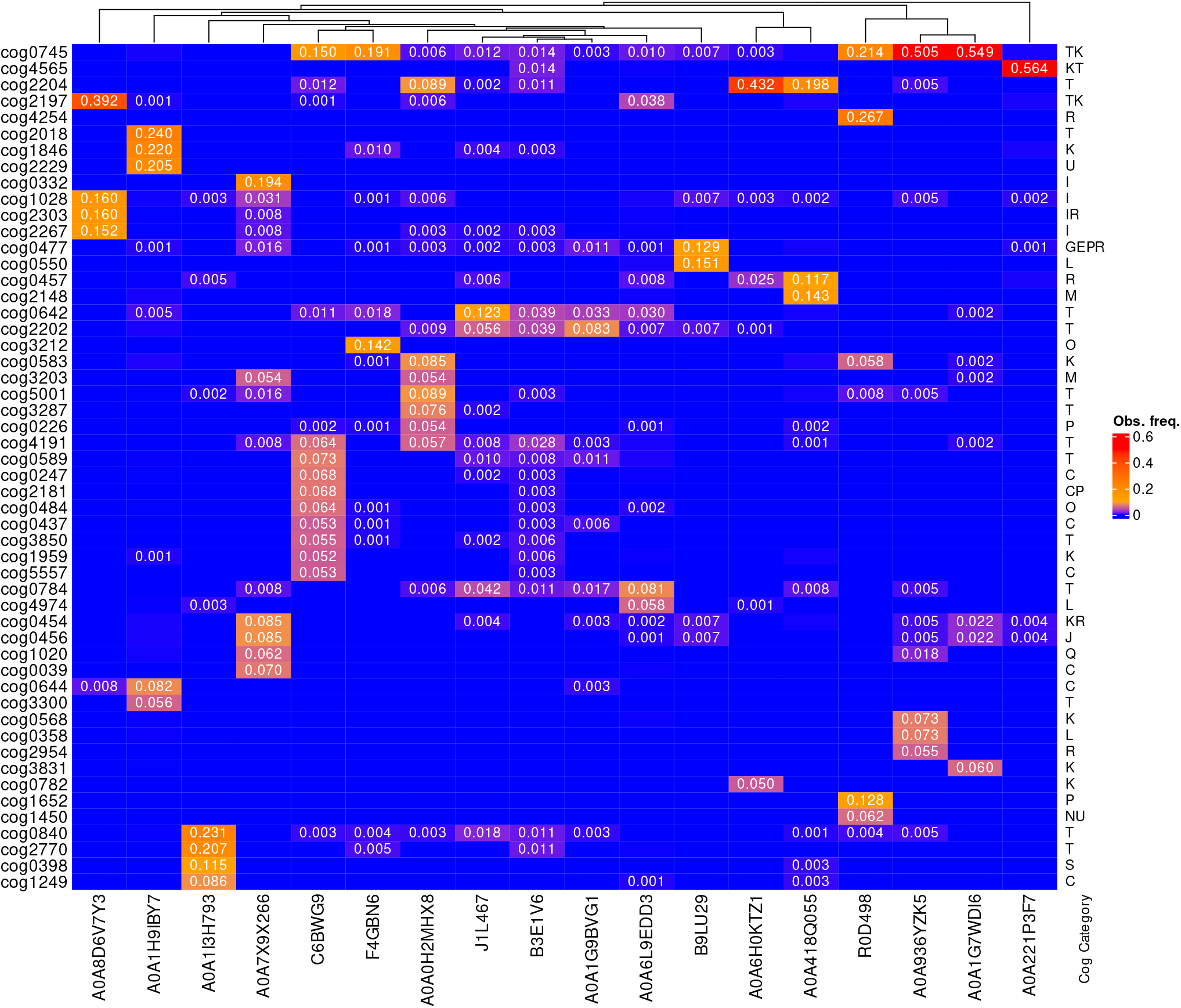
Heatmap of the observed frequency of COGs in the genomic context of the 18 profiles. Rows correspond to COGs and columns to profiles. Annotations on the right indicate the COG category of the corresponding COG on the left. Only COGs with an observed frequency *>* 0.05 in at least one profile are shown. Values below 0. 001 are not reported. The dendogram at the top represent a hierarchical clustering between the profiles based on COG assignment.

Only six COGs were shared across multiple profiles: COG0745 (OmpR response regulator family), COG2204 (NtrC response regulator family), COG2202 (PAS domain), COG0583 (LysR family transcriptional regulators), COG3203 (OmpC/OmpF/PhoE outer membrane proteins) and COG4191 (histidine kinases regulating C4-dicarboxylate transport). As shown in Figure 12, no COG was markedly over-represented in any genomic context; the most frequently occurring COGs appeared in at most 20% of the genomic neighborhoods across all profiles. This suggests that the profiles are not associated with specific regulatory activities and do not share any regulatory function in a statistically meaningful way.

### 3.5 Negative dataset

As a sanity check, we aligned our profiles against a set of sequences differing from standard HKs based on their EC number and domain architecture. Our negative dataset comprised 439 sequences, three of which matched two of our profiles: A0A1G9BVG1 and J1L467, with one and two matches, respectively. Table 3 lists these matches along with their associated UniProt annotations and EC numbers.

**Table 3:** Matches between the negative dataset and our profiles.

| Profile | UniProt ID | UniProt annotation | EC number |
| --- | --- | --- | --- |
| A0A1G9BVG1 | Q0W3A5 | Signal transduction serine/threonine kinase | 2.7.11.1 |
| J1L467 | A0A2V2N3N6 | histidine kinase | 2.7.7.72 |
|  | A0A2V2ND00 | histidine kinase | 2.7.7.72 |

The protein matching A0A1G9BVG1 is annotated as a serine/threonine kinase (STK) in UniProt, consistent with its EC number. This protein is 1 827 amino acids long and carries STK-related domains in its N-terminal region and HK-related domains (including HATPase and PS50109) in its C-terminal region. Its domain annotation suggests both histidine kinase and serine/threonine kinase activity, making its true function ambiguous. Furthermore, this protein should not have been included in the negative dataset, given its association with the PROSITE profile PS50109 in UniProt (see section 2.4). The fact that this match was not detected by InterProScan highlights the annotation inconsistencies that can arise between standard tools and databases.

The two proteins matching J1L467 are annotated as HKs in InterPro, whereas their EC numbers classify them as CCA tRNA nucleotidyltransferases. Both proteins carry multiple HK-related domains, including PAS and REC domains, identified across several databases (Pfam, SMART, PROSITE, CDD, and CATH-Gene3D). In UniProt, both are also associated with the PROSITE profile PS50109. Taken together, these observations suggest that the EC number assignment is misleading in these cases and that these proteins are likely iHKs that should not have been included in the negative dataset. Overall, these findings indicate that the three matches identified in the negative dataset correspond to sequences that incorrectly passed the filtering steps. They should therefore not be considered as evidence that profiles A0A1G9BVG1 and J1L467 lack specificity for HisKA domains.

### 3.6 iHKs found in models organisms

Using our profiles, we searched for iHKs in 41 organisms of known interest (see Supplementary Table S7). We identified 112 matches in 22 organisms, spanning 16 of our 18 profiles (the two exceptions being A0A1G9BVG1 and A0A936YZK5). The proteome files analyzed may contain sequences annotated as “hypothetical proteins” as a result of the automated NCBI annotation pipeline; however, only one of the 112 matches involved such a sequence. Some matched proteins carried HK-related terms in their annotation (e.g. “two-component sensor” or “sensor histidine kinase”), while others had more generic annotations such as “PAS domain-containing protein” or “ATP-binding protein”. For the latter, the identification of an HisKA-like domain could help clarify their function. Table 4 summarizes the organisms and the number of matches obtained. RefSeq IDs of the matched proteins and the profiles they matched are provided in Supplementary Tables S8 and S9.

**Table 4:**
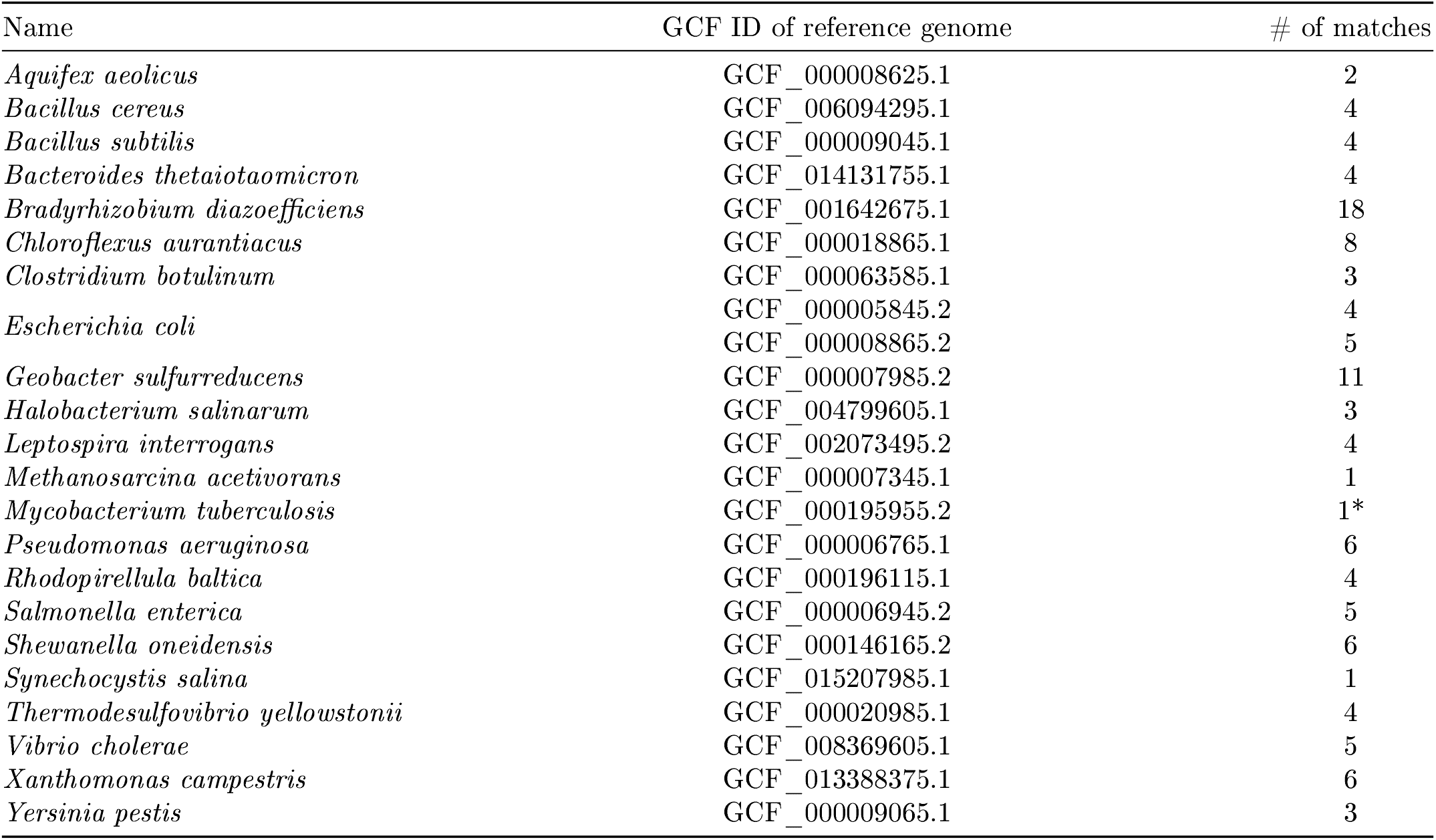
Names of the 22 well-studied organisms in which iHKs matching our profiles were identified. The middle column gives the GCF accession of their reference genome. The right column gives the number of iHKs identified in each organism. An asterisk (^*^) denotes a hypothetical protein.

## 4 Discussion

Using a large-scale computational analysis of more than 800 000 iHK sequences, we identified 18 HisKA-like HMM profiles, each available in both SEED and FULL versions. These profiles, named after representative UniProt entries, were subjected to a series of sequence-, structure-, and context-based analyses to evaluate their validity as HisKA domains and their relevance relative to existing annotation resources.

(Willett and Kirby, 2012) showed that the kinase and phosphatase activities of CrdS in *Myzococcus xanthus* are linked to specific conserved residues in the HisKA domain. They demonstrated that an acidic residue (E/D) adjacent to the histidine residue is required for kinase activity, while phosphatase activity appears to depend on the presence of a threonine or asparagine residue (T/N) four amino acids downstream of the conserved histidine. However, these findings may not generalize to all HisKA domain types, as some lack these conserved residues (e.g. PF06580 lacks an acidic residue after the conserved histidine, and PF19191 lacks a conserved threonine/asparagine residue) . Nevertheless, 14 of our profiles (J1L467, A0A221P3F7, A0A1I3H793, F4GBN6, A0A1G7WDI6, R0D498, A0A6H0KTZ1, A0A7X9X266, A0A8D6V7Y3, A0A936YZK5, A0A418Q055, A0A0H2MHX8, B9LU29 and A0A1G9BVG1) possess these conserved residues, further supporting their probable kinase and phosphatase activities.

Taxonomic analysis of the iHKs present in our FULL profiles revealed that all profiles are specific to bacteria and archaea. No matches were found with plant or fungal sequences, at least among the iHKs analyzed in this study. It is not impossible that our profiles could identify sequences from eukaryotic genomes not included here. This contrasts with PF00512, the most general HisKA profile in Pfam, which spans both prokaryotic and eukaryotic HKs. All other HisKA profiles in Pfam are found exclusively in prokaryotes, though none are archaea-specific (PF07568 and PF19191 match both bacteria and archaea, while all others are bacteria-only).

We found that several of our profiles aligned well with SwissProt proteins for which the phosphorylatable histidine residue has been experimentally or confidently annotated, and that our profiles correctly identified the same histidine residue in these sequences. However, some of these proteins are also annotated with other known profiles (SSF47384, SSF55890 and SM00388) that likewise identify the phosphorylatable histidine residue. By construction, our profiles should not exhibit similarities to known profiles, given the numerous sequence filtering steps applied (section 2.1). These conflicting annotations arise from differences in the thresholds used by tools and databases to assign matches: in our case, InterProScan, which was used to filter sequences, did not report these matches, whereas UniProt did. Regardless, for most sequences affected by these conflicting annotations, our profiles achieved higher alignment scores than those from SuperFamily, SMART, and Pfam. There were five exceptions, involving sequences that aligned more closely with SSF55890 than with our profile A0A221P3F7. To facilitate future annotations, each of our profiles was assigned a threshold analogous to the gathering threshold (GA) used in Pfam profiles.

One of our profiles (A0A0H2MHX8) also matched the atypical histidine kinase Lpl0330. By comparing the conserved histidine residue identified by profile A0A0H2MHX8 with that characterized by Levet-Paulo et al., we found that A0A0H2MHX8 was not centered on the correct histidine residue. When aligning the complete sequences from the SEED MSA of A0A0H2MHX8, we discovered two columns with conserved histidine residues in the HisKA-like domain region. The initial version of A0A0H2MHX8 was centered on the more C-terminal histidine residue, which was in fact located too close to the HATPase domain. We therefore rebuilt A0A0H2MHX8 based on the conserved histidine residue of Lpl0330. Profile A0A0H2MHX8 was the only case in which multiple fully conserved histidine residues were present in the region of a potential HisKA domain.

Using AlphaFold2-Multimer, we predicted the homodimeric 3D structure of each profile representative. As stated previously, the dimer models produced by AlphaFold2-Multimer should be taken cautiously, as the relationships between the different subunits are not determined with enough confidence. However, each of our HisKA-like profiles are found in regions having distinct folds and well defined in the dimers. In all predicted structures, two *α*-helices were found at the location of the HisKA-like domain, consistent with the known structural features of HisKA domains described in the literature. Furthermore, for all profiles except A0A1H9IBY7, the conserved histidine residue was correctly positioned within the structure, i.e. at the midpoint of the first *α*-helix. The structural classification of the regions aligning with our profiles yielded classifications similar to those of the HisKA domains in the ECOD, CATH and TED databases. For some profiles a complete structural classification was not possible, as it often end with sequence-based associations. These new profiles, if integrated in these databases, could help increase classification coverage. We performed pairwise structural alignments with curated reference HisKA domain structures and reference HisKA-HATPase complexes using TM-Align. They revealed notable similarities, further supporting the validity of the majority of profiles. We also studied the local structural environment of the predicted phosphorylatable histidine residues in each representative’s HisKA-like domain and saw that they share structural features. Moreover, when there is more than one histidine residue in a HisKA-like domain, we saw that it was possible to distinguish the predicted phosphorylatable histidine residue based on its position in the *α*-helices and its solvent-accessible surface areas. Profile A0A1H9IBY7 stood out numerous time as an outsider in many analysis, raising doubts about its HisKA-like characteristics. Indeed, the structural features of its predicted phosphorylatable histidine residue are not as expected. It is located in the second *α*-helix and its SASA value is significantly lower than those of the other phosphorylatable histidine residues. Unlike A0A0H2MHX8, no additional conserved histidine column was identified near the one initially selected. In the absence of *in vitro* or *in vitro* data, it is most prudent to conclude that this profile does not correspond to an HisKA domain. While it could represent an atypical HisKA variant, the strong structural conservation of the HisKA fold across all known HK proteins argues against this hypothesis.

Using eggNOG-mapper, we analyzed the genomic context of genes present in our SEED alignments to determine whether they are associated with regulatory activities, as is typically the case for HK genes. We retrieved COG categories and assignments, where available, for all genes identified as neighbors of our iHKs. There are 26 COG categories and over 5 000 COG assignments; the latter are more specific and often correspond to discrete protein domains, while COG categories provide broader functional classifications. We conducted two separate analyses: one focused on COG categories and one on individual COG assignments. COG categories in the genomic contexts indicate that genes associated with our profiles tend to occur near genes involved in regulation and signal transduction (categories K and T). Profile B9LU29 was the only exception, showing low representation of both categories T and K. Category S (“Function unknown”) was also strongly represented across most genomic contexts, which is unsurprising given that this study focuses on poorly characterized proteins. Interestingly, some profiles appear to share similarities at the level of COG categories: for example, profiles A0A8D6V7Y3 and A0A7X9X266 both show a high proportion of category I (“Lipid transport and metabolism”) . While these patterns could suggest a functional grouping of profiles based on their genomic context, this was not supported by COG assignment analysis. COG assignments did not systematically mirror COG categories, with some genes carrying a COG category annotation but a COG assignment from a different functional class. Ultimately, only six COG assignments were found at significant frequency across more than one profile, and none were present at levels sufficient to indicate specific shared regulatory activities. These results suggest that the profiles identified in this study are associated with genes found in highly diverse genomic contexts, consistent with involvement in a broad range of regulatory pathways.

Profile specificity was further evaluated by screening against non-HK proteins. Only three matches were detected across all profiles, one of which corresponded to a protein clearly annotated as non-HK in UniProt; however, this protein carried multiple HK-related domains in its C-terminal region. Overall, no profile showed systematic or convincing association with proteins clearly unrelated to HKs, indicating a high degree of specificity. Searching for iHKs in well-studied prokaryotic organisms using the profiles described in this study yielded 112 matches across 22 model organisms, spanning 16 of the 18 profiles, with only one match falling on a hypothetical protein. This demonstrates that the proposed profiles can contribute to the characterization of regulatory pathways in well-studied organisms.

## 5 Conclusion

Although 18 HisKA-like profiles were identified, additional variants were likely missed due to the genome quality and clustering criteria applied during profile construction. The absence of eukaryotic profiles most likely reflects the low abundance of iHKs in these genomes rather than a limitation of the methodology. Beyond amino acid composition, it is not yet possible to determine precisely how and why these profiles differ from known HisKA profiles. Structural analysis did not show divergent structural features from known HisKA domains. In the end, several profiles — most notably A0A0H2MHX8 — are supported by converging sequence, structural, and contextual evidence. Profile A0A1H9IBY7, however, should be interpreted with caution, as its predicted 3D structure deviates from the canonical HisKA fold. In the absence of experimental validation, the robustness of individual profiles remains variable.

Taken together, these results demonstrate that the proposed HMM profiles constitute a refined and complementary resource to existing HisKA annotations, well suited for integration into large-scale genome annotation pipelines. We believe these profiles could facilitate the discovery of novel HKs in prokaryotic regulatory pathways and, given their application to well-studied organisms, could prove valuable to researchers across a range of fields, including health and ecology. Moreover, the methodology developed in this study, documented in full in the GitLab repository, can be applied more broadly to other domain types defined by a conserved residue or sequence motif.

## 6 Declarations

### 6.1 Ethics approval and consent to participate

Not applicable

### 6.2 Consent for publication

Not applicable

### 6.3 Availability of data and materials

The sequences of iHKs analyzed during this study, the SEED and FULL profiles and the 3D structure of each profile representative are available in the public repository at https://doi.org/10.57745/Y9N0P9, through the service Recherche Data Gouv. The scripts, command lines and complete methodology used in this study are detailed in the GitLab repository at https://gitlab.n2p3.Mr/louison.silly/hiska-like-domain-characterization.

## 6.5 Funding

This work was supported by the doctoral school Evolution, Ecosystems, Microbiology and Modelisation at University Claude Bernard + Lyon 1.

### 6.6 Authors’ contributions

LS and PO did all the developments, analyzes and testing required for this study. PO and GP supervised the analyzes performed by LS. LS wrote the manuscript and drew the figures which were completed and corrected by GP and PO.

## 6.7 Acknowledgments

Structure predictions and some MSAs were computed on the Jean Zay supercomputer facility at the Institute for Development and Resources in Scientific Computing in Paris-Saclay.

---

**Table S1:** List of known HisKA domains in Pfam, SMART, SuperFamily, CDD and CATH-Gene3D.

| Entry ID | Database |
| --- | --- |
| PF00512 | Pfam |
| PF07568 |  |
| PF07730 |  |
| PF07536 |  |
| PF06580 |  |
| PF19191 |  |
| PF14689 |  |
| PF22561 |  |
| SM00388 | SMART |
| SM00911 |  |
| SSF47384 | SuperFamily |
| SSF55890 |  |
| cd00082 | CDD |
| G3DSA:1.20.5.1930 | CATH-Gene3D |
| G3DSA:1.10.287.130 |  |
| PS50109 | PROSITE |

**Table S2:**
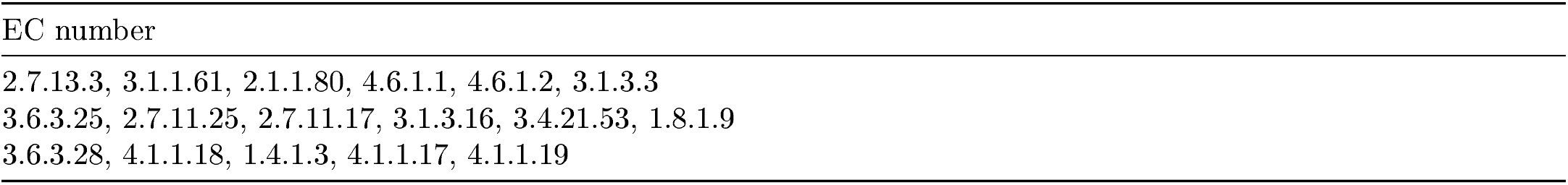
List of EC number to search for after the annotation of iHK with eggNOG-mapper.

**Table S3:**
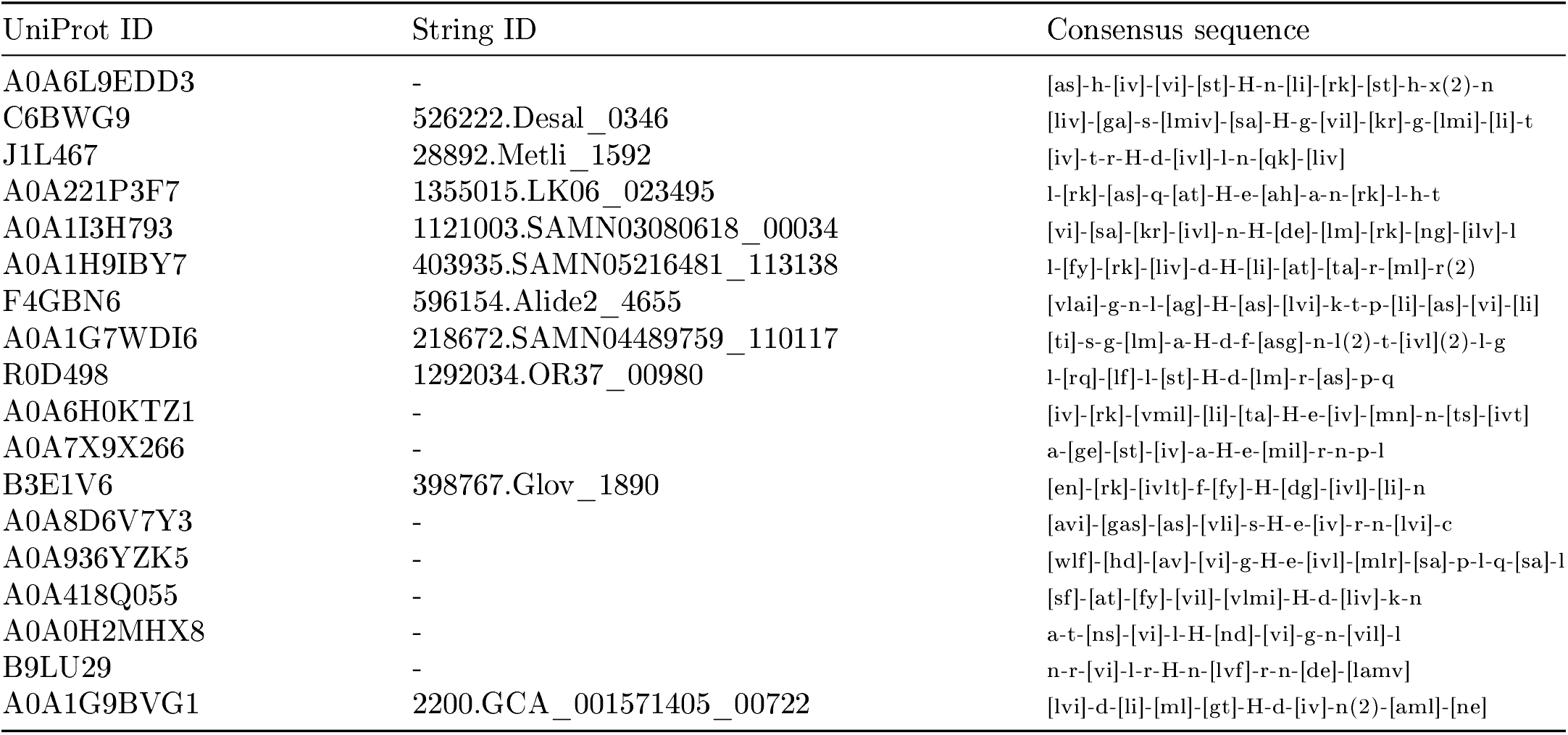
UniProt ID, STRING ID (where available), and consensus sequence surrounding the conserved histidine residue for each of the 18 profiles. In the consensus sequences, all residues are shown in lowercase except for the conserved histidine. When a position contains multiple residues, they are ordered by decreasing probability of occurrence. The degenerate residue X is used for positions where no residue reaches a probability ≥ 0.1. When one or more residues are repeated consecutively, they are preceded by (*n*), where *n* denotes the number of repetitions.

**Table S4:** List of the 27 SwissProt sequences that align with three of our profiles. **SwissProt annotation** indicates the position of the phosphohistidine as manually annotated in SwissProt. **Profile annotation** indicates the position of the putative phosphohistidine identified by the profile listed in the **Profile** column. For all sequences, the same histidine residue is identified by both SwissProt and our profiles.

| Sequence ID (UniProt) | Profile | SwissProt<br>annotation | Profile<br>annotation |
| --- | --- | --- | --- |
| Q9I0I2 | F4GBN6 | 239 | 239 |
| Q8X739 | F4GBN6 | 277 | 277 |
| Q83RR1 | F4GBN6 | 277 | 277 |
| Q8FIB8 | F4GBN6 | 277 | 277 |
| P23837 | F4GBN6 | 277 | 277 |
| D0ZV89 | F4GBN6 | 277 | 277 |
| E1WFA0 | F4GBN6 | 277 | 277 |
| F5ZP94 | F4GBN6 | 277 | 277 |
| P0DM80 | F4GBN6 | 277 | 277 |
| Q57QC4 | F4GBN6 | 277 | 277 |
| Q5PMJ0 | F4GBN6 | 277 | 277 |
| Q8Z7H3 | F4GBN6 | 277 | 277 |
| Q9RC53 | A0A221P3F7 | 342 | 342 |
| P52687 | A0A221P3F7 | 350 | 350 |
| O34427 | A0A221P3F7 | 339 | 339 |
| P77510 | A0A221P3F7 | 347 | 347 |
| P96601 | A0A221P3F7 | 336 | 336 |
| Q9K997 | A0A221P3F7 | 339 | 339 |
| O05250 | A0A221P3F7 | 338 | 338 |
| P0AEC8 | A0A221P3F7 | 349 | 349 |
| P0AEC9 | A0A221P3F7 | 349 | 349 |
| P59341 | A0A221P3F7 | 349 | 349 |
| P59340 | A0A221P3F7 | 349 | 349 |
| P0C5S6 | A0A7X9X266 | 471 | 471 |
| A7MRY4 | A0A7X9X266 | 471 | 471 |
| Q9KM66 | A0A7X9X266 | 194 | 193 |
| A7N6S2 | A0A7X9X266 | 190 | 190 |

**Table S5:** Alignment scores of 27 SwissProt sequences against profiles from SuperFamily, SMART, Pfam, and our in-house profiles. Proteins are grouped by the custom profile they match (F4GBN6, A0A221P3F7, or A0A7X9X266). Some proteins share high sequence similarity and therefore yield identical alignment scores. In most cases, these proteins align more closely with our profiles than with those from SuperFamily, SMART, and Pfam.

| UniProt ID | Profiles alignment scores |  |  |
| --- | --- | --- | --- |
|  | SSF47384 |  | F4GBN6 |
| Q9I0I2 | 23.67 |  | **78.78** |
| Q8X739 | 20.34 |  | **37.27** |
| Q83RR1 | 20.36 |  | **37.03** |
| Q8FIB8 | 20.36 |  | **37.03** |
| P23837 | 20.36 |  | **37.03** |
| D0ZV89 | 19.83 |  | **35.86** |
| E1WFA0 | 19.83 |  | **35.86** |
| F5ZP94 | 19.83 |  | **35.86** |
| P0DM80 | 19.83 |  | **35.86** |
| Q57QC4 | 19.83 |  | **35.86** |
| Q5PMJ0 | 19.83 |  | **35.86** |
| Q8Z7H3 | 19.83 |  | **35.86** |
|  | SSF55890 | PF14689 | A0A221P3F7 |
| Q9RC53 | 46.05 | 37.67 | **72.77** |
| P52687 | 24.01 | 18.30 | **67.85** |
| O34427 | 35.57 | 33.13 | **62.32** |
| P77510 | 30.60 | 26.66 | **62.53** |
| P96601 | 41.59 | 36.65 | **61.49** |
| Q9K997 | 38.81 | 34.27 | **59.71** |
| O05250 | **55.28** | 41.72 | 46.38 |
| P0AEC8 | **57.65** | 33.53 | 41.18 |
| P0AEC9 | **57.65** | 33.53 | 41.18 |
| P59341 | **57.65** | 33.53 | 41.18 |
| P59340 | **57.65** | 33.53 | 41.18 |
|  | SSF47384 | SM00388 | A0A7X9X266 |
| P0C5S6 | 22.69 | 28.01 | **34.44** |
| A7MRY4 | 22.69 | 28.01 | **34.44** |
| Q9KM66 | 21.45 | 21.41 | **35.24** |
| A7N6S2 | 21.76 | 22.00 | **31.43** |

**Table S6:** Average pLDDT of the HisKA-like domains in the monomeric counterparts of the representatives structures. Last column on the right display the pLDDT score for the conserved histidine residue in the domains.

| Representative's ID | Average pLDDT of the HisKA-like domain | pLDDT of the conserved his residue |
| --- | --- | --- |
| A0A0H2MHX8 | 86.93 | 66.62 |
| A0A1G9BVG1 | 80.90 | 80.94 |
| A0A1I3H793 | 63.80 | 63.38 |
| A0A418Q055 | 72.95 | 70.0 |
| A0A6L9EDD3 | 80.71 | 79.12 |
| A0A8D6V7Y3 | 70.55 | 62.69 |
| B3E1V6 | 84.49 | 72.19 |
| C6BWG9 | 79.57 | 70.12 |
| J1L467 | 85.13 | 77.88 |
| R0D498 | 77.60 | 75.12 |
| A0A1G7WDI6 | 64.08 | 57.31 |
| A0A1H9IBY7 | 73.17 | 58.22 |
| A0A221P3F7 | 73.0 | 70.44 |
| A0A6H0KTZ1 | 86.10 | 85.38 |
| A0A7X9X266 | 68.31 | 64.12 |
| A0A936YZK5 | 70.06 | 69.19 |
| B9LU29 | 82.01 | 78.19 |
| F4GBN6 | 82.62 | 81.94 |

**Table S7:**
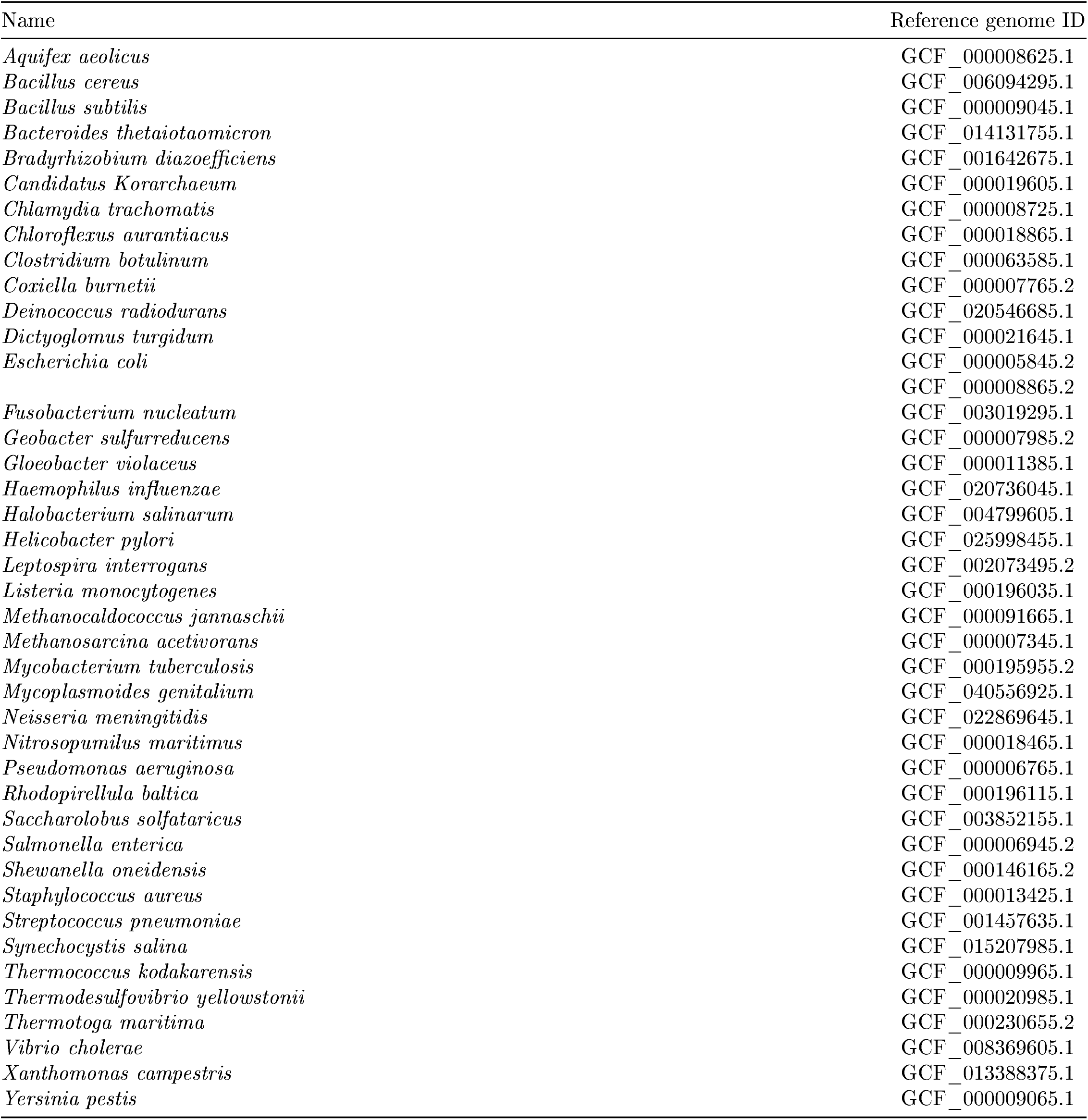
List of the 41 model organisms in which we searched for iHKs using our in-house HisKA-like profiles. GCF accession numbers correspond to the reference genomes of each organism in the NCBI database. *Escherichia coli* is the only entry with two reference genomes.

**Table S8:** RefSeq IDs of the iHKs found in 22 model organisms, and the profile they matched.

| Organisms name | RefSeq IDs of the matches | Name of the profiles |
| --- | --- | --- |
| Aquifex aeolicus | WP_010880154.1<br>WP_010880552.1 | F4GBN6<br>A0A6H0KTZ1 |
| Bacillus cereus | WP_033685471.1<br>WP_000749654.1;WP_002025277.1;WP_000746421.1 | A0A6H0KTZ1<br>A0A221P3F7 |
| Bacillus subtilis | NP_388639.1;NP_388326.1;NP_391030.1<br>NP_389282.1 | A0A221P3F7 |
| Bacteroides<br>thetaiotaomicron | WP_011107401.1;WP_011107493.1;WP_011107842.1<br>WP_011107804.1 | A0A6H0KTZ1 |
| Bradyrhizobium<br>diazoefficiens | WP_011085906.1<br>WP_038967572.1<br>WP_011086340.1<br>WP_028175987.1<br>WP_028171469.1;WP_011090098.1;WP_162494129.1<br>WP_011084968.1;WP_011088025.1;WP_011090708.1<br>WP_011083217.1;WP_011088423.1;WP_011083347.1<br>WP_038966842.1;WP_011089455.1;WP_014497709.1<br>WP_038967507.1;WP_011088552.1 | F4GBN6<br>A0A6H0KTZ1<br>A0A7X9X266<br>A0A1I3H793<br>A0A1G7WDI6 |
| Chloroflexus<br>aurantiacus | WP_012257996.1;WP_012259340.1;WP_012256371.1<br>WP_242605105.1;WP_242604947.1;WP_012257124.1<br>WP_012256308.1;WP_012256764.1 | A0A1G7WDI6 |
| Clostridium<br>botulinum | WP_011986463.1<br>WP_011948042.1<br>WP_012099532.1 | A0A6L9EDD3<br>A0A221P3F7<br>R0D498 |
| Escherichia coli<br>(GCF_000005845.2) | NP_415647.1<br>NP_416126.1<br>NP_415152.1;NP_418549.1 | F4GBN6<br>A0A7X9X266<br>A0A221P3F7 |
| Escherichia coli<br>(GCF_000008865.2) | NP_309628.1<br>NP_313101.1<br>NP_310342.2<br>NP_308685.1;NP_313134.1 | F4GBN6<br>A0A1G7WDI6<br>A0A7X9X266<br>A0A221P3F7 |
| Geobacter<br>sulfurreducens | WP_010942586.1<br>WP_010942829.1<br>WP_010943621.1<br>WP_235044997.1;WP_010943038.1;WP_010941506.1<br>WP_010942299.1;WP_235045002.1;WP_010943442.1<br>WP_010942202.1;WP_010941951.1 | A0A418Q055<br>B3E1V6<br>R0D498<br>A0A1G7WDI6 |
| Halobacterium<br>salinarum | WP_244288991.1;WP_244288990.1;WP_136361441.1 | B9LU29 |
| Leptospira<br>interrogans | WP_000997518.1<br>WP_000490514.1<br>WP_000581337.1;WP_001288616.1 | A0A6H0KTZ1<br>A0A1G7WDI6<br>A0A6L9EDD3 |

**Table S9:**
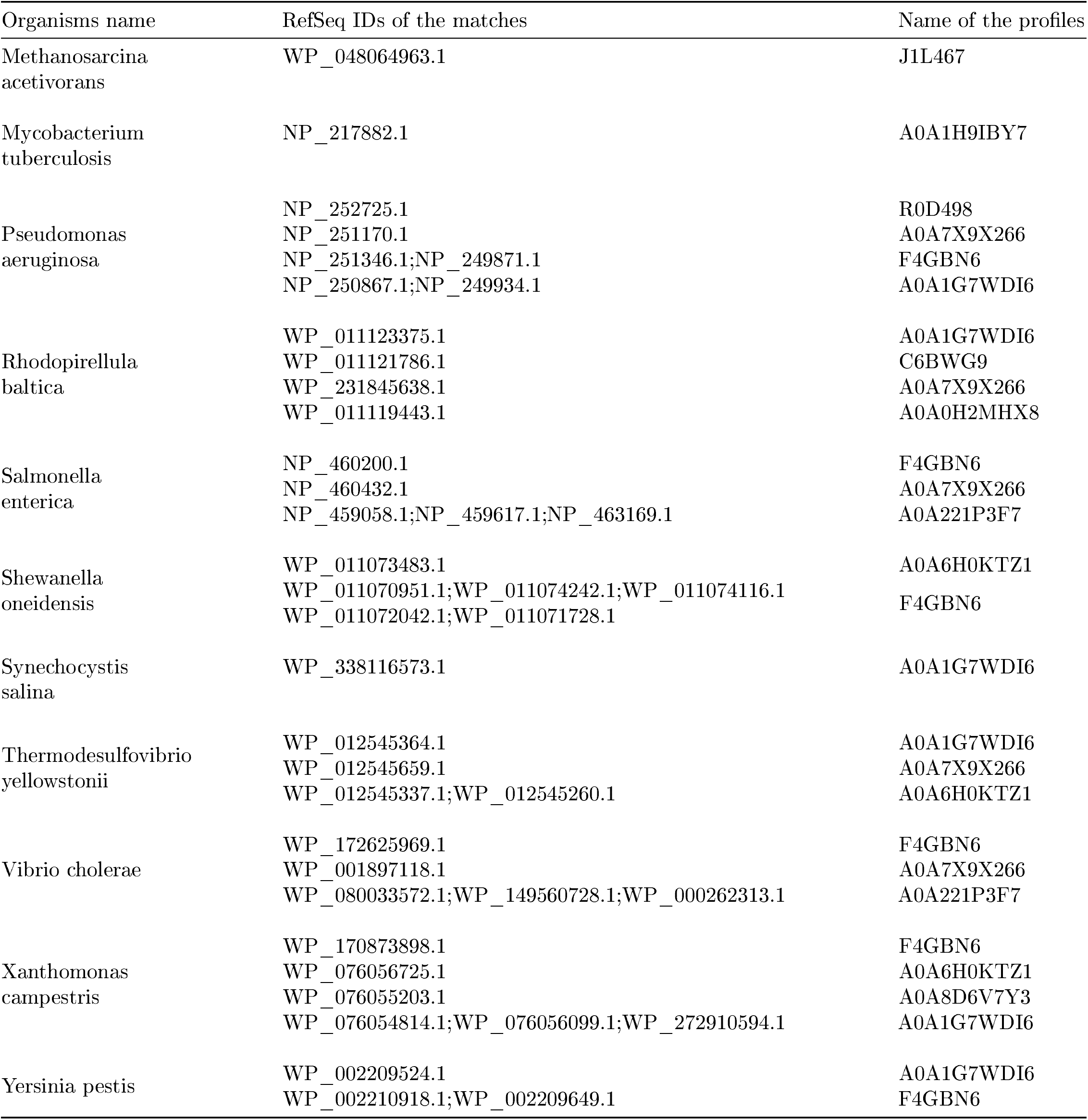
Following of table S8.

**Figure S1:**
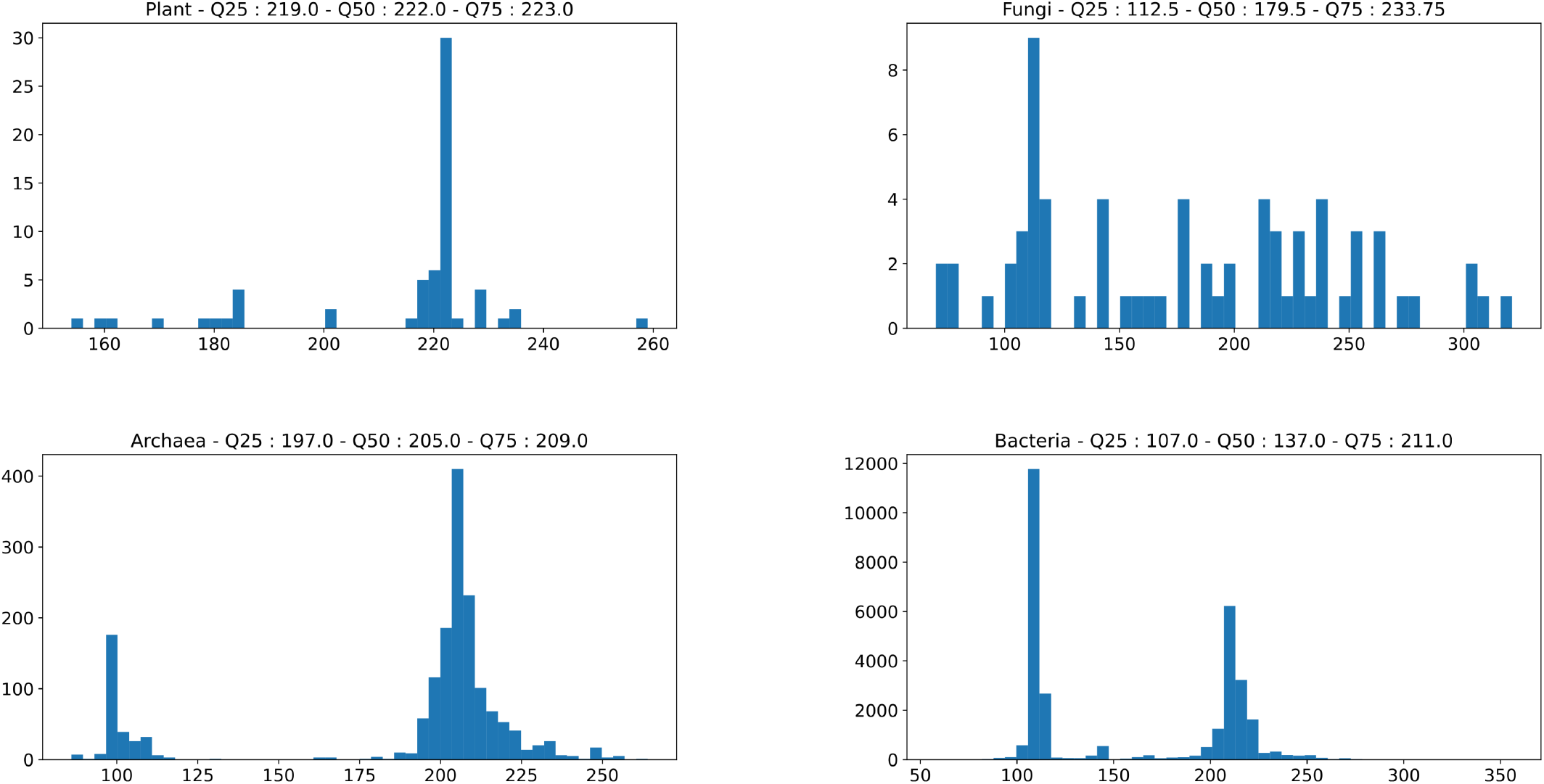
Distribution of match lengths against the PROSITE profile PS50109 for each group of iHK sequences, after filtering. Quartiles (0.25, 0.50, and 0.75) are indicated in each plot title.

**Figure S2:**
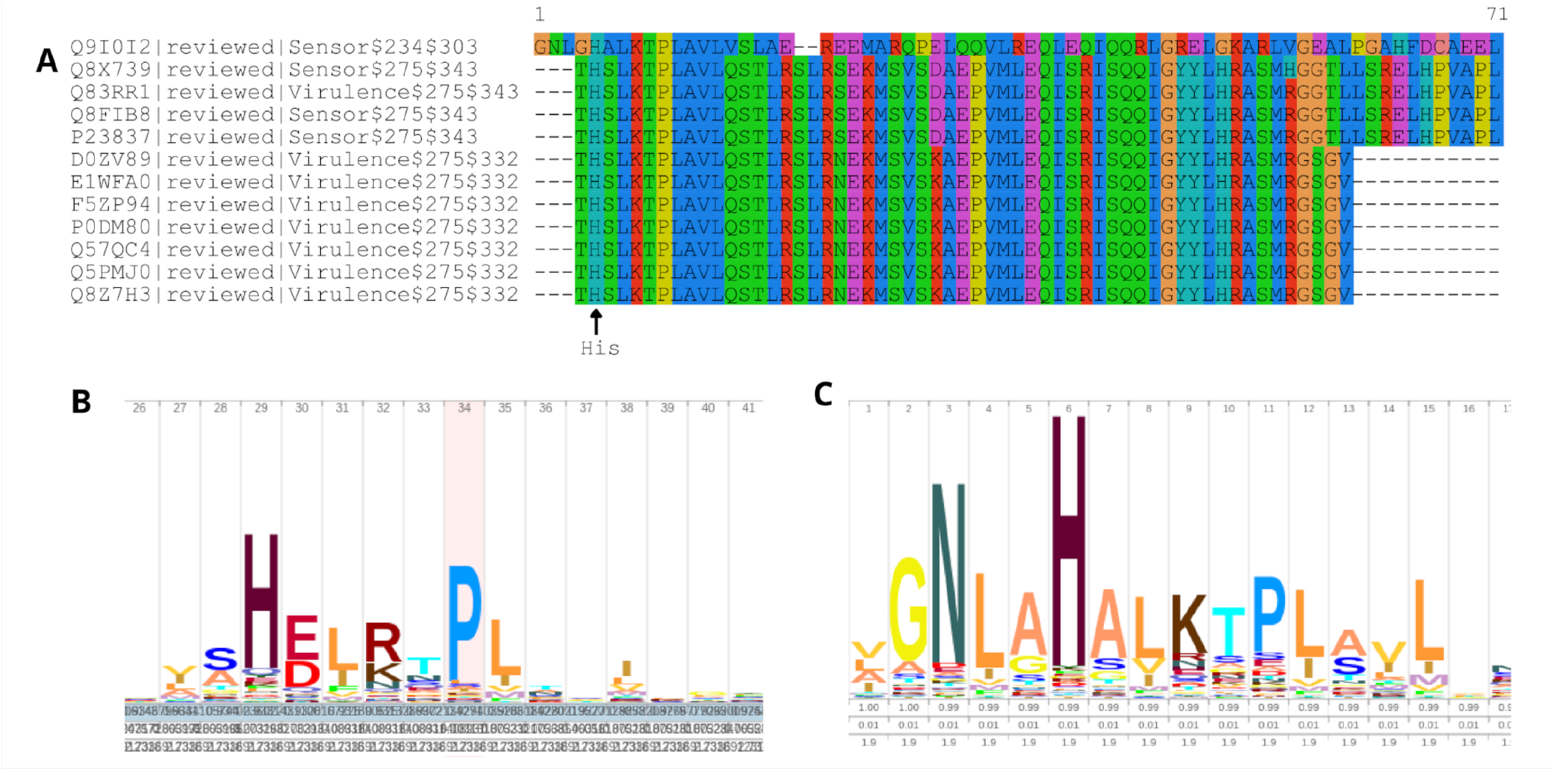
**A**: Alignment, displayed with SeaView, of 12 SwissProt sequences matching both our profile F4GBN6 and the SuperFamily profile SSF47384. The conserved histidine residue column is at position 5. **B**: Sequence logo of the H-Box from SSF47384, retrieved from the SSF47384 entry page on the InterPro website. **C**: Sequence logo of the H-Box from F4GBN6, generated using Skylign. It more closely reflects the alignment than the sequence logo of SSF47384, particularly for the first amino acids following the conserved histidine.

**Figure S3:**
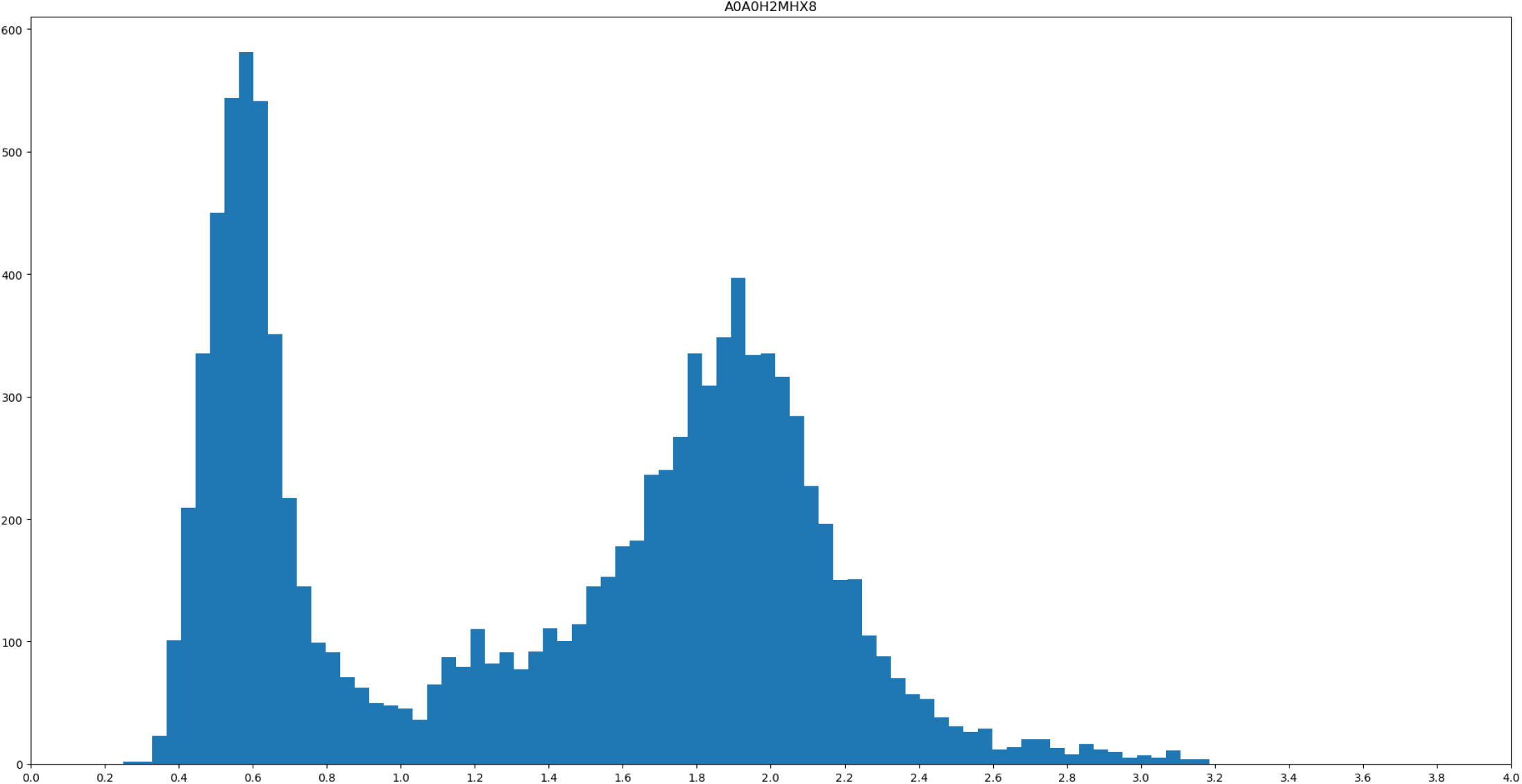
Sequence logo of profile A0A0H2MHX8 – neighborhood of the conserved histidine residue (−5/+19).

**Figure S4:**
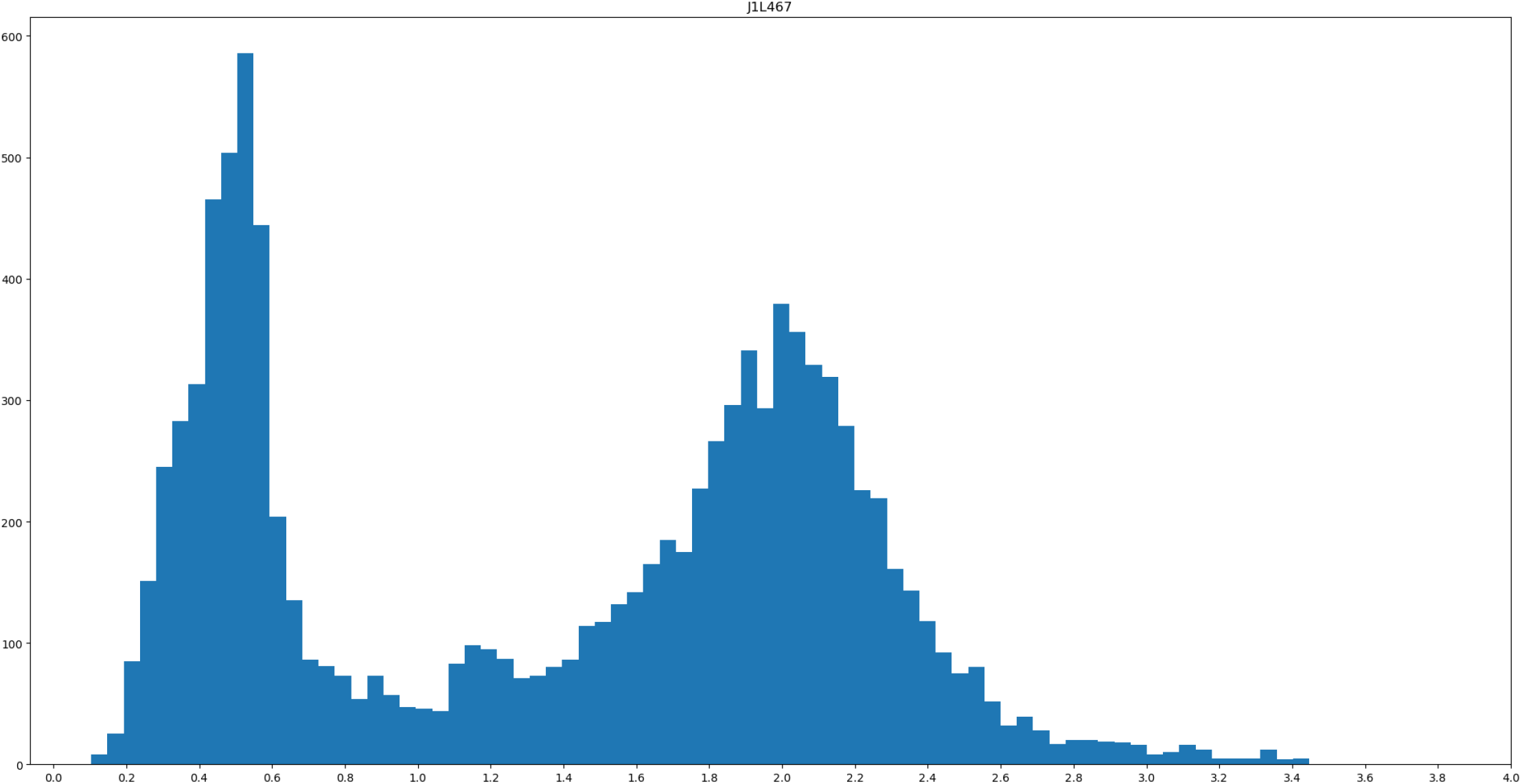
Sequence logo of profile J1L467 – neighborhood of the conserved histidine residue (−5/+19).

**Figure S5:**
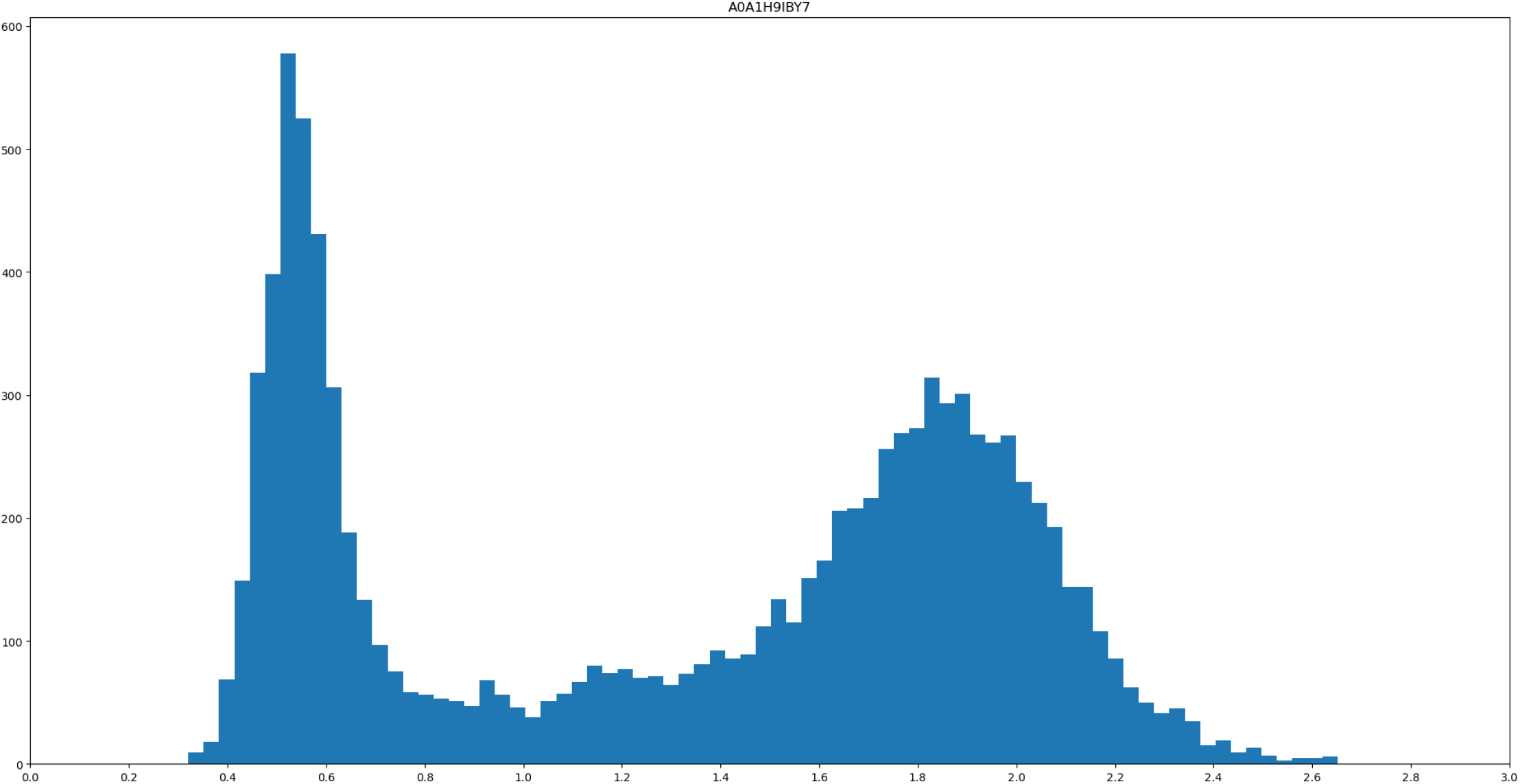
Sequence logo of profile A0A1H9IBY7 – neighborhood of the conserved histidine residue (−5/+19).

**Figure S6:**
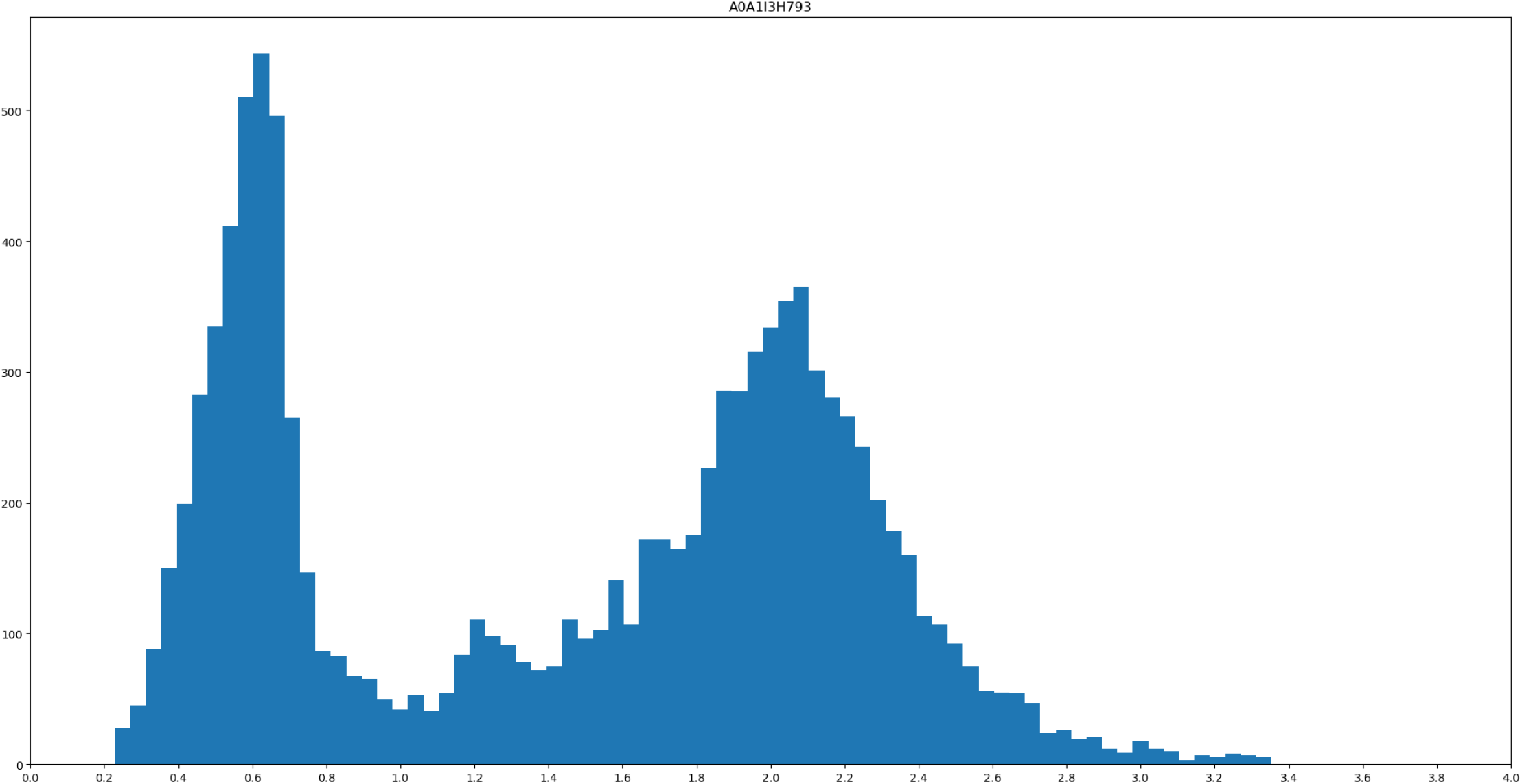
Sequence logo of profile A0A1I3H793 – neighborhood of the conserved histidine residue (−5/+19).

**Figure S7:**
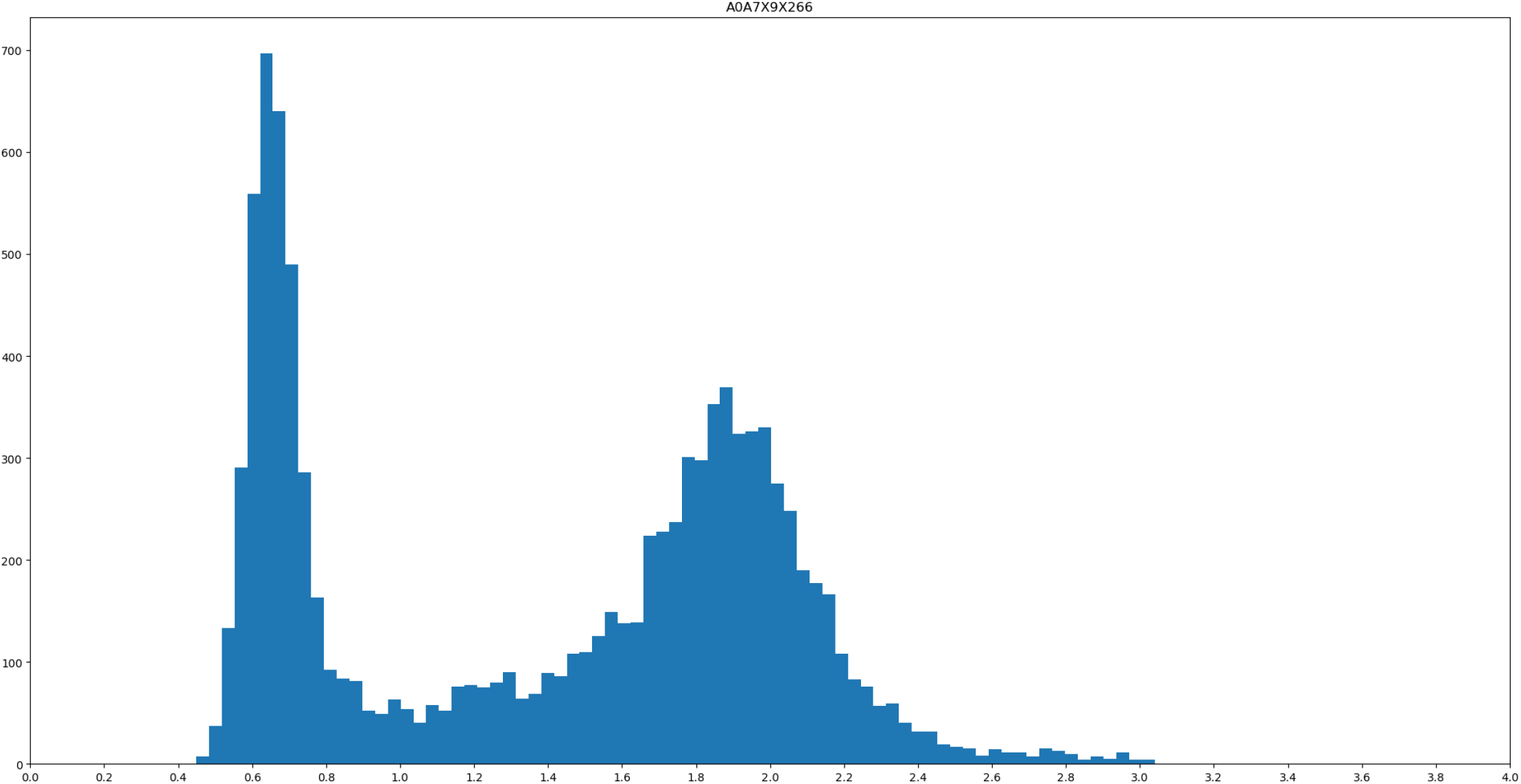
Sequence logo of profile A0A7X9X266 – neighborhood of the conserved histidine residue (−5/+19).

**Figure S8:**
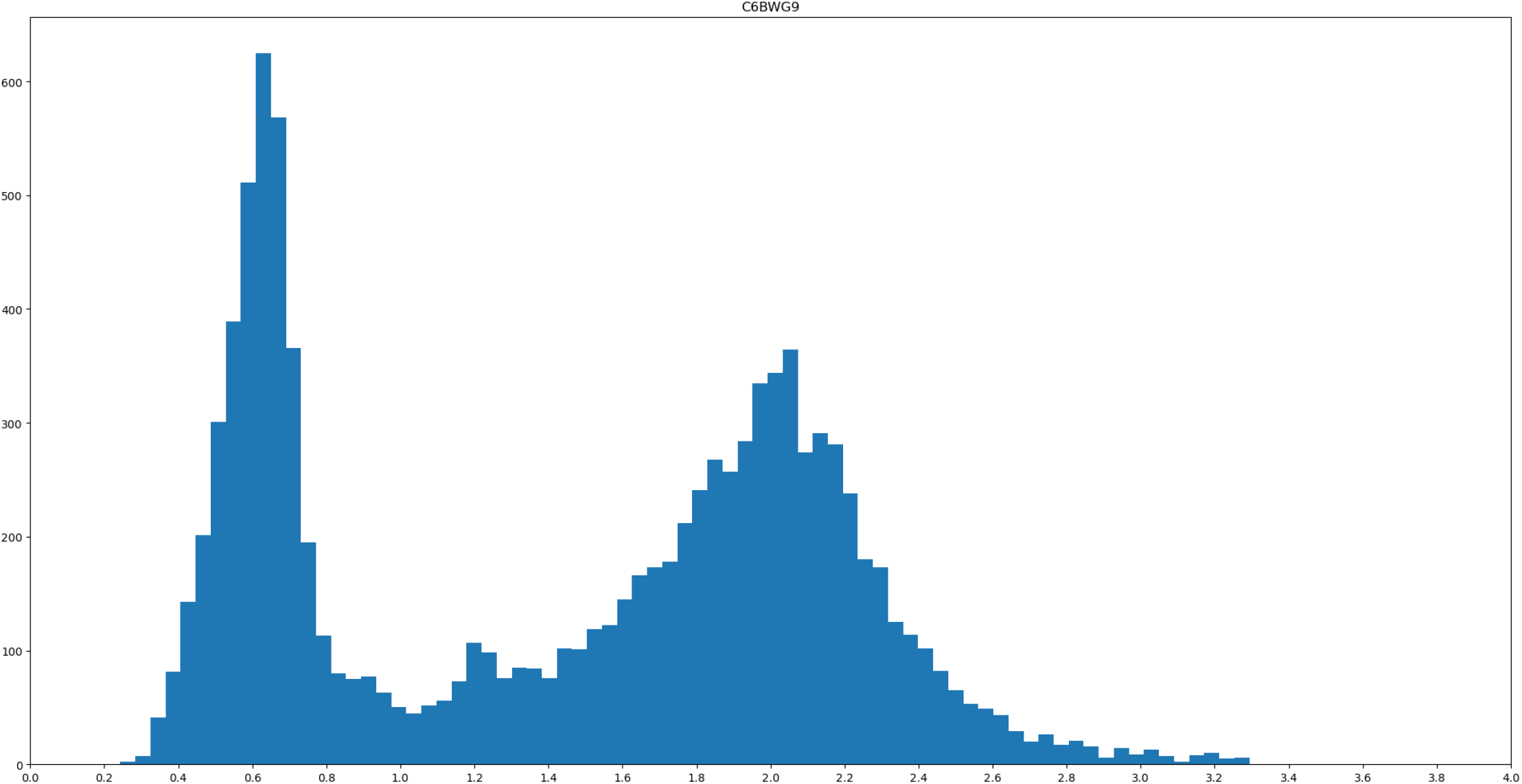
Sequence logo of profile C6BWG9 – neighborhood of the conserved histidine residue (−5/+19).

**Figure S9:**
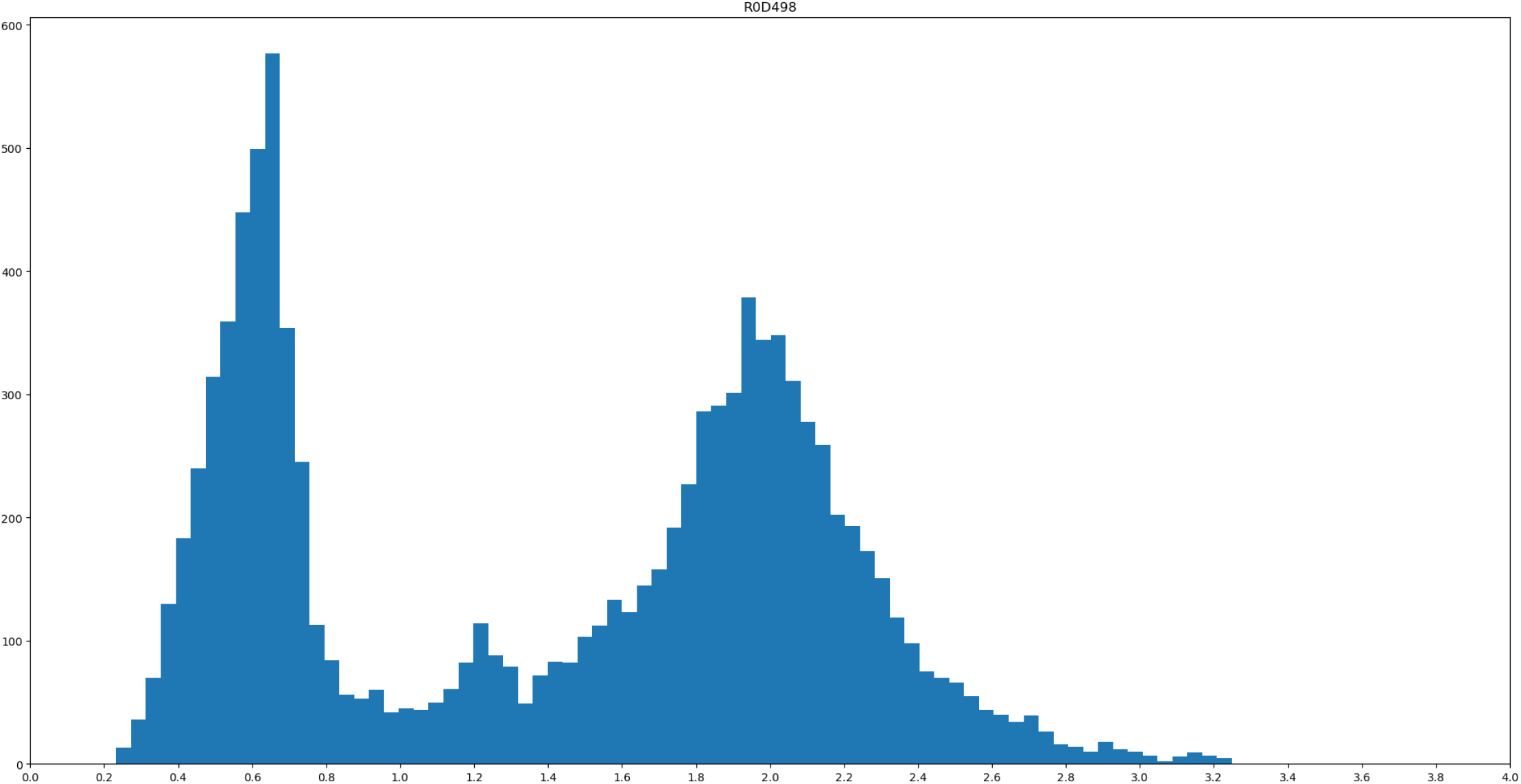
Sequence logo of profile R0D498 – neighborhood of the conserved histidine residue (−5/+19).

**Figure S10:**
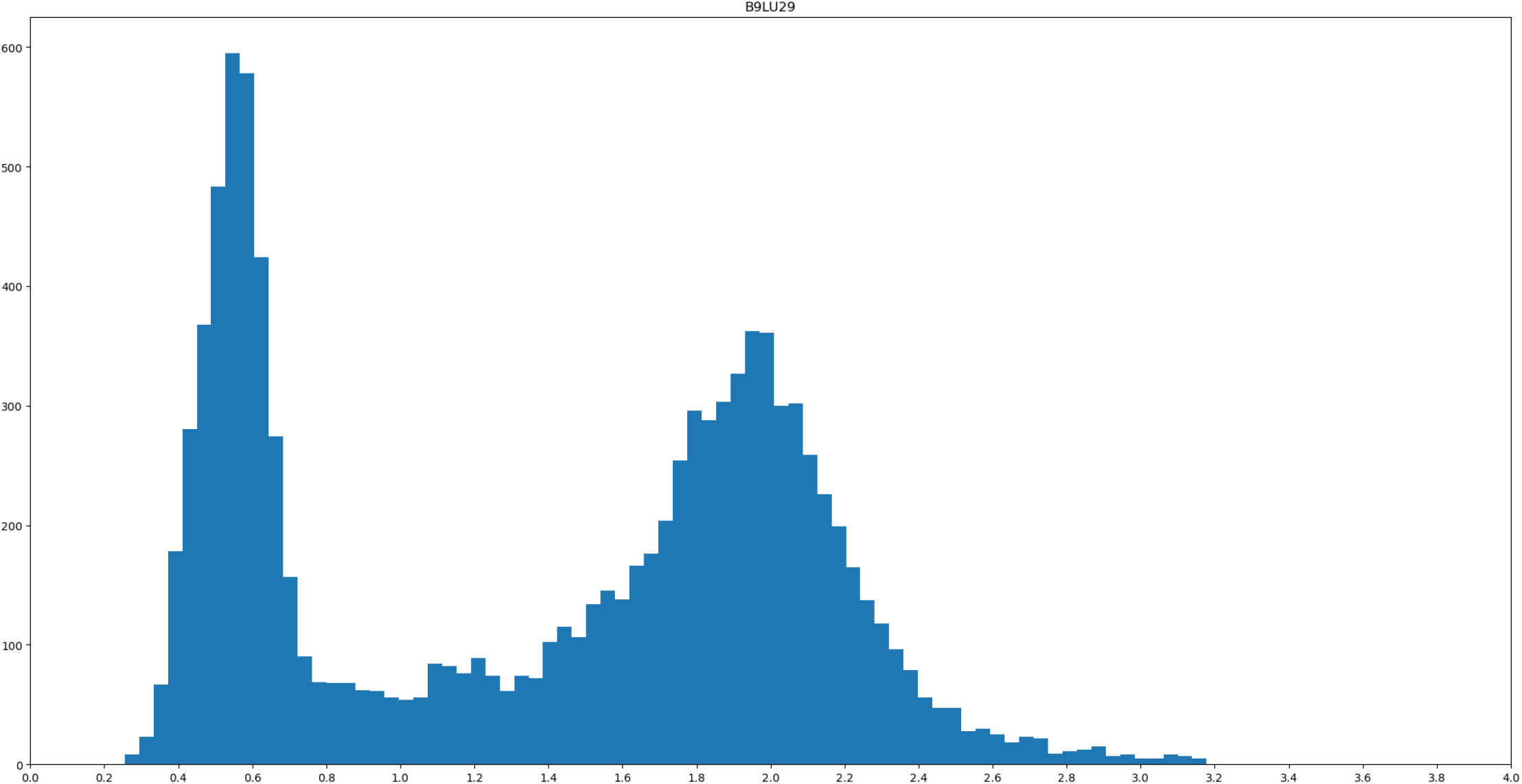
Sequence logo of profile B9LU29 – neighborhood of the conserved histidine residue (−5/+19).

**Figure S11:**
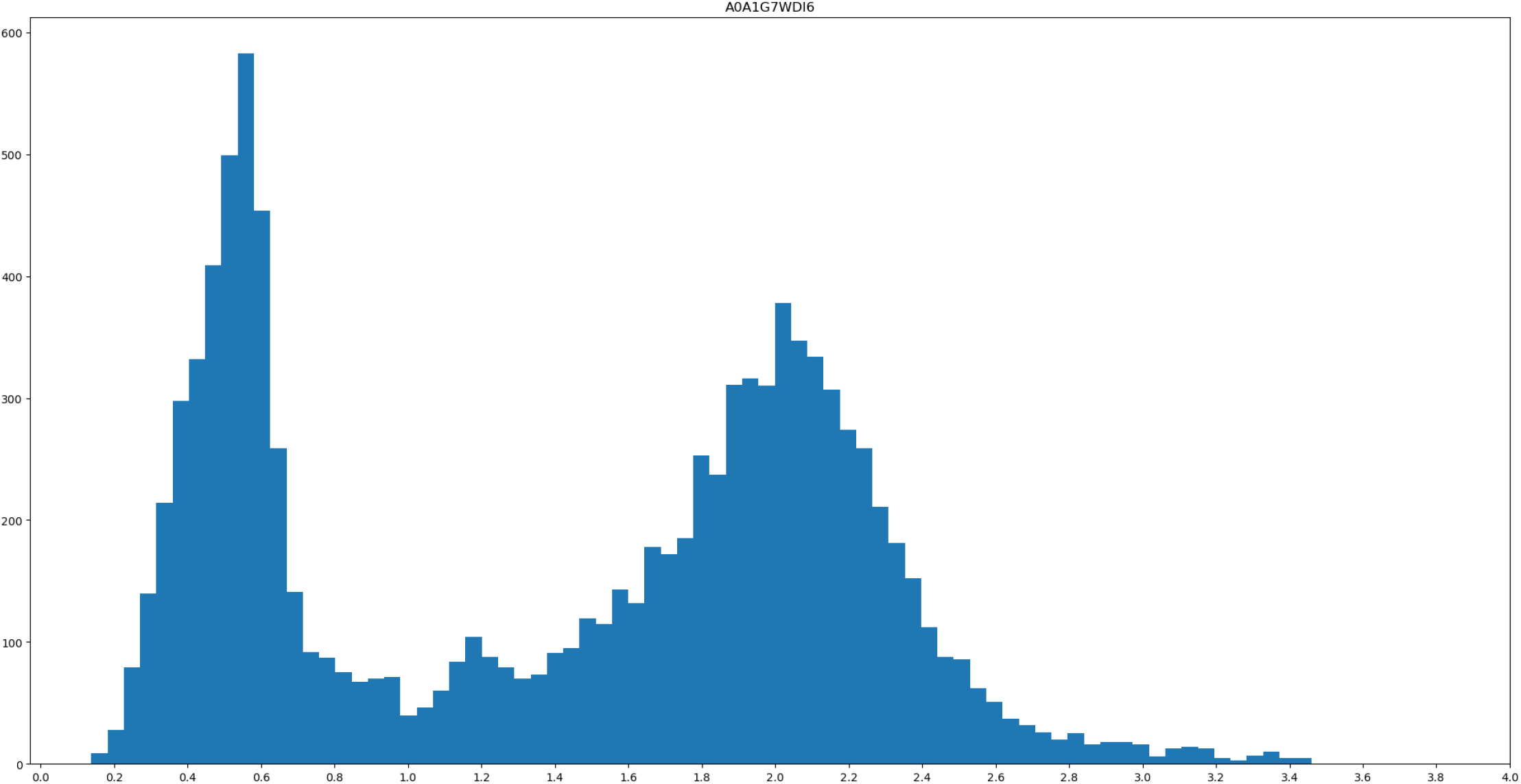
Sequence logo of profile A0A1G7WDI6 – neighborhood of the conserved histidine residue (−5/+19).

**Figure S12:**
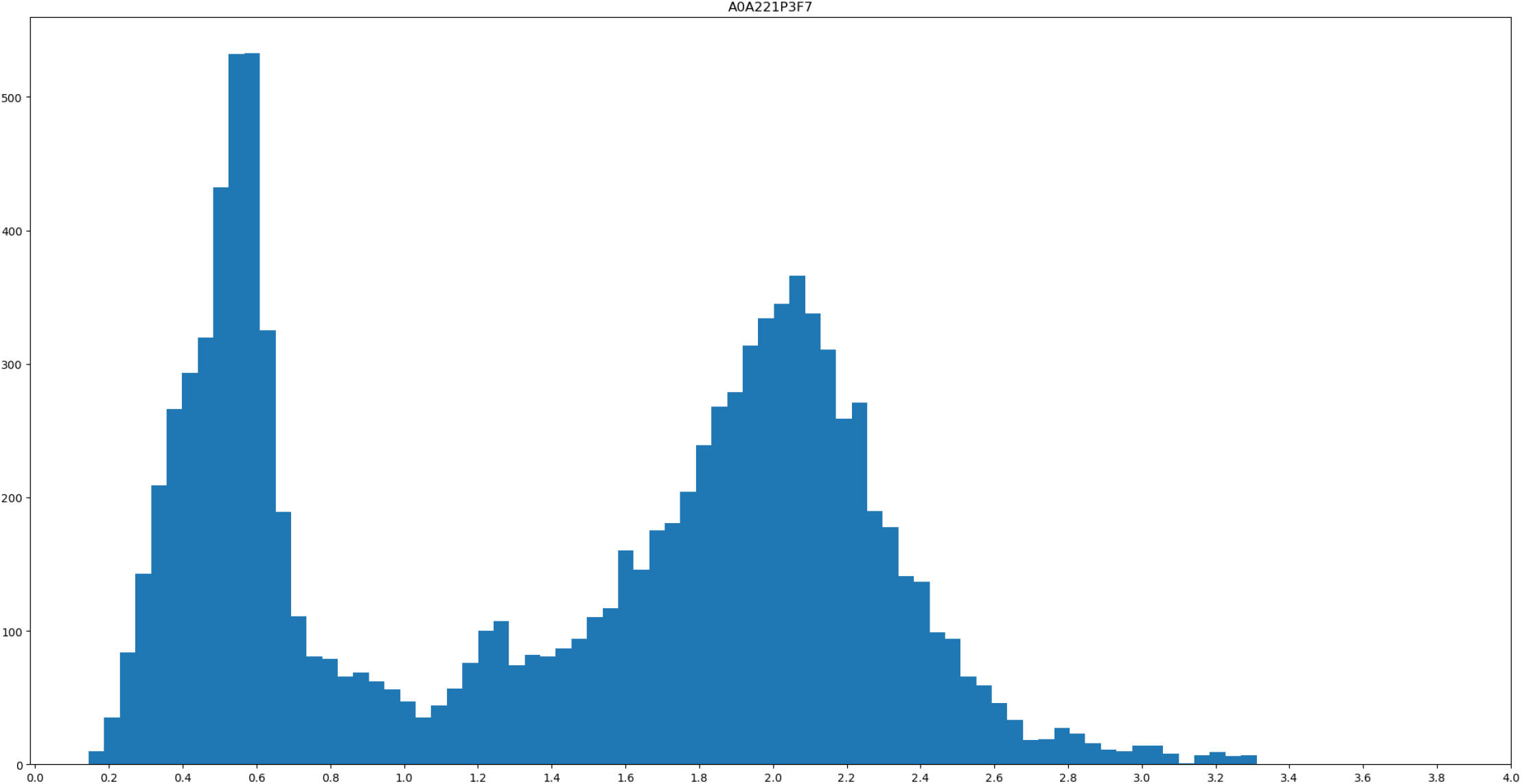
Sequence logo of profile A0A221P3F7 – neighborhood of the conserved histidine residue (−5/+19).

**Figure S13:**
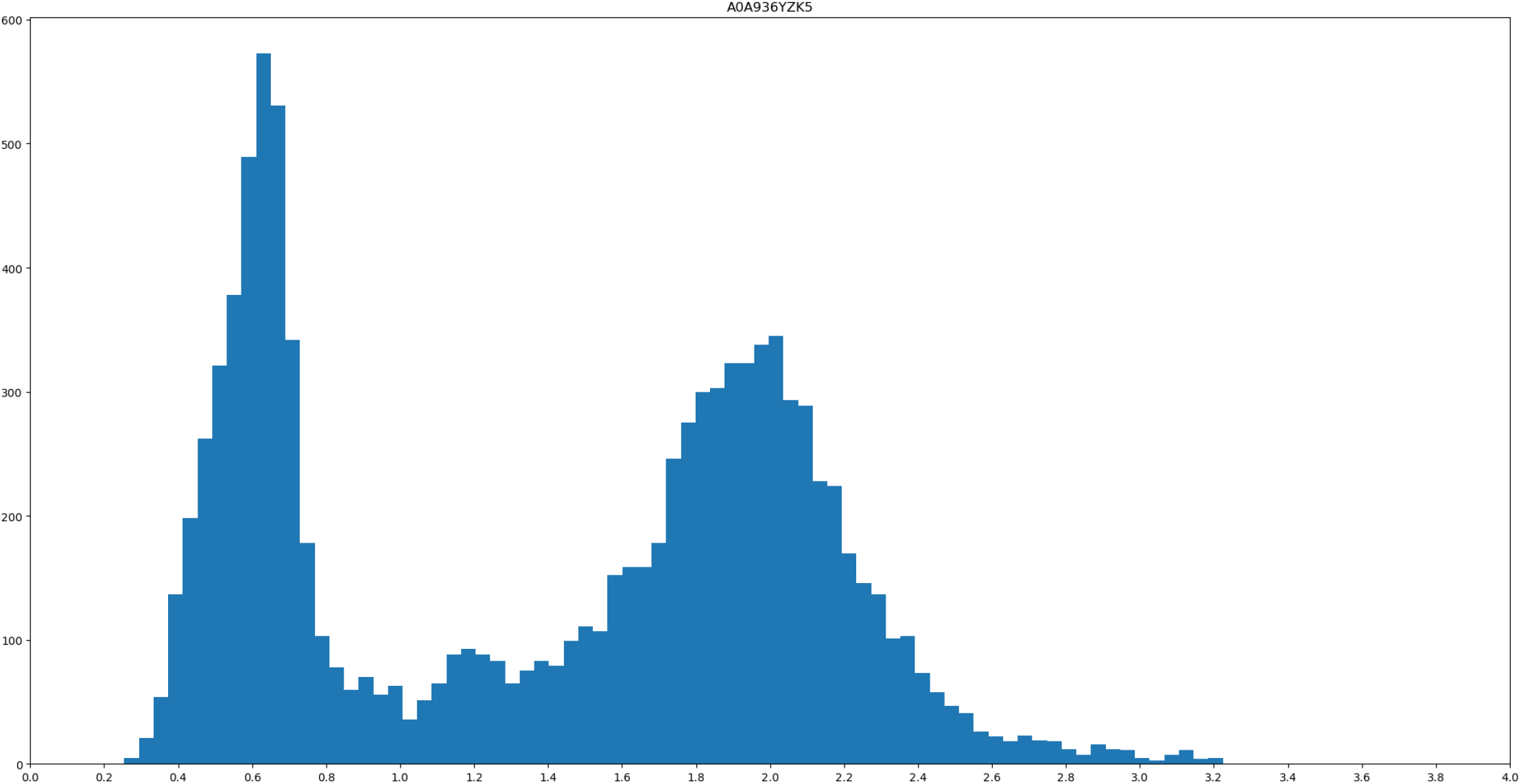
Sequence logo of profile A0A936YZK5 – neighborhood of the conserved histidine residue (−5/+19).

**Figure S14:**
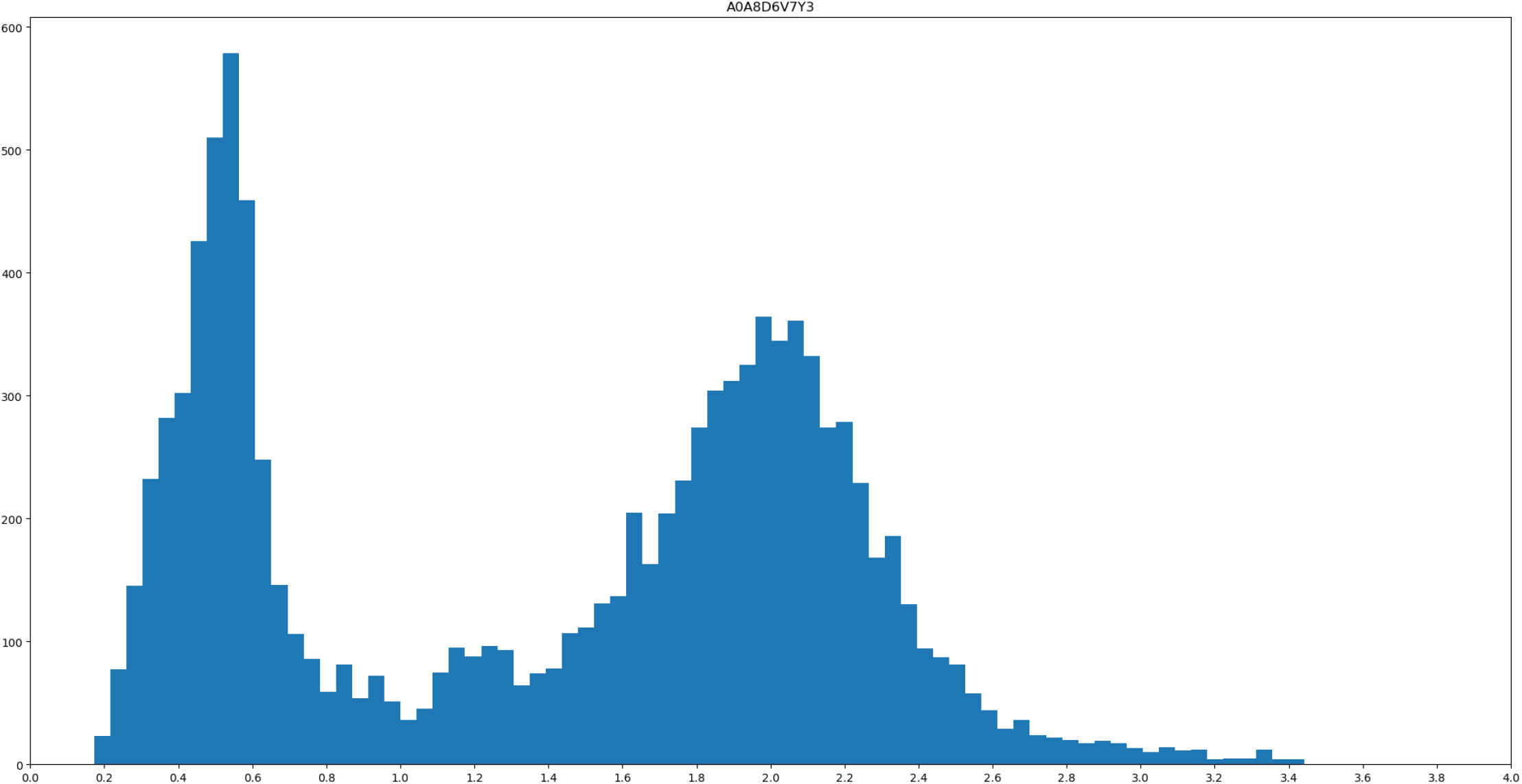
Sequence logo of profile A0A8D6V7Y3 – neighborhood of the conserved histidine residue (−5/+19).

**Figure S15:**
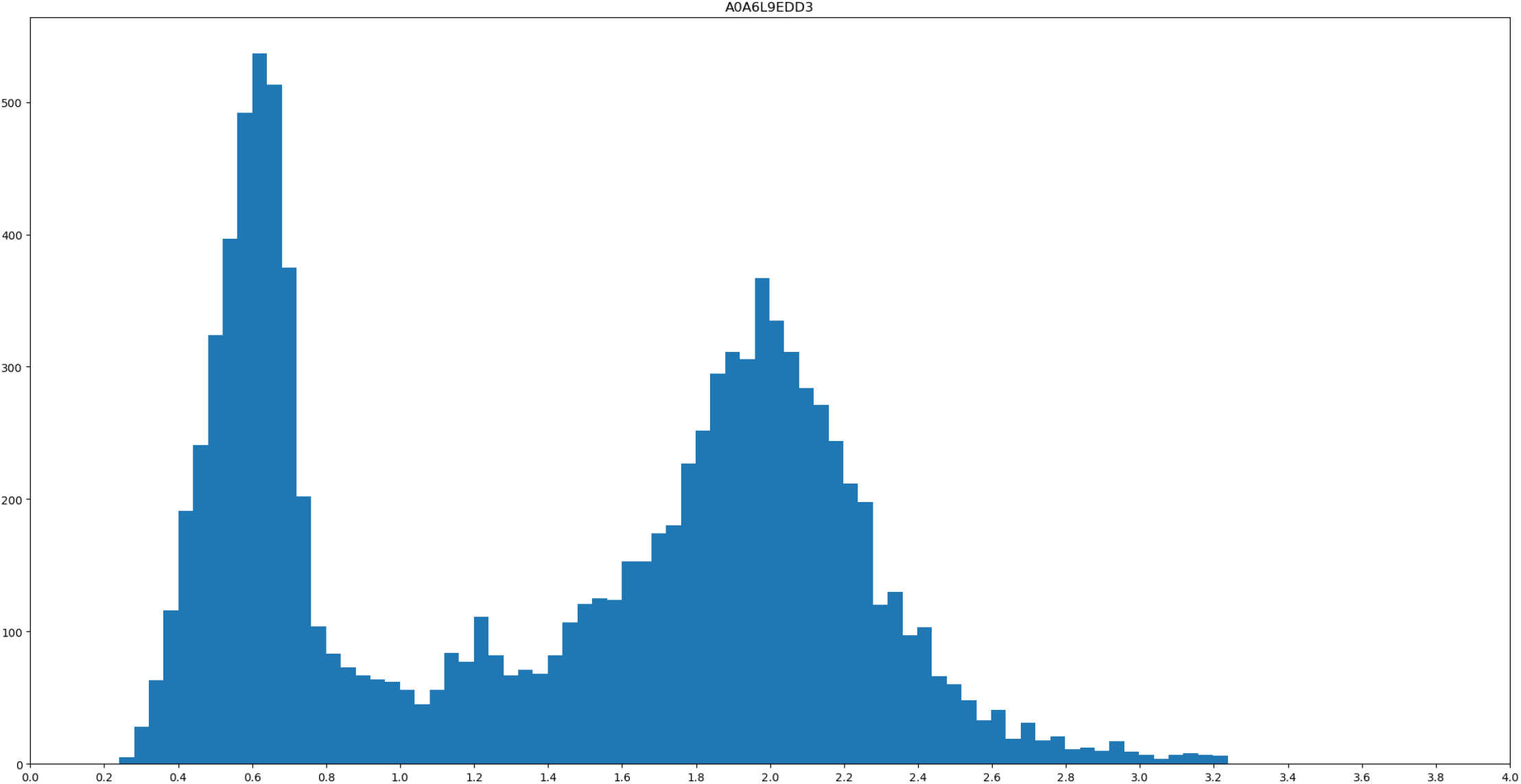
Sequence logo of profile A0A6L9EDD3 – neighborhood of the conserved histidine residue (−5/+19).

**Figure S16:**
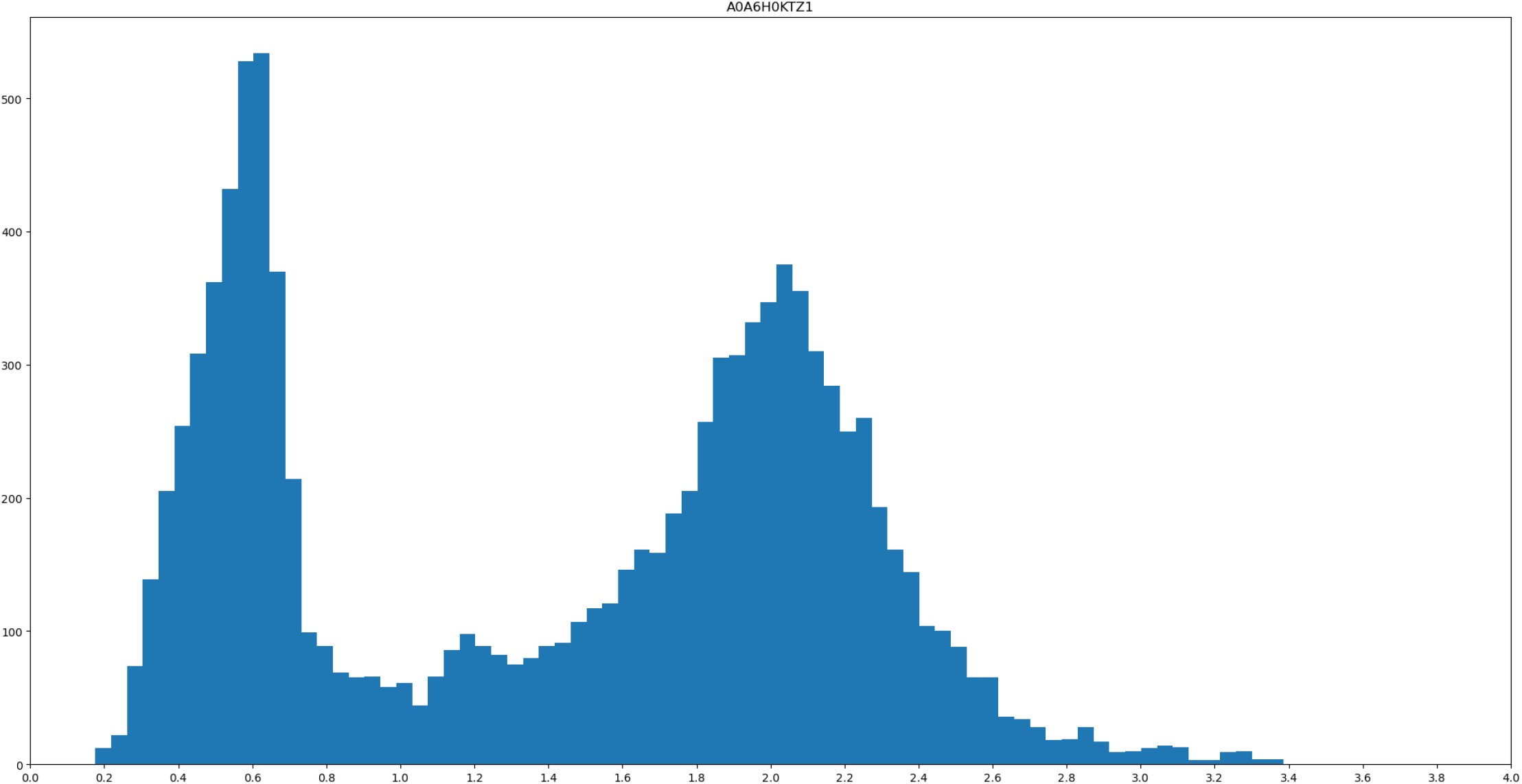
Sequence logo of profile A0A6H0KTZ1 – neighborhood of the conserved histidine residue (−5/+19).

**Figure S17:**
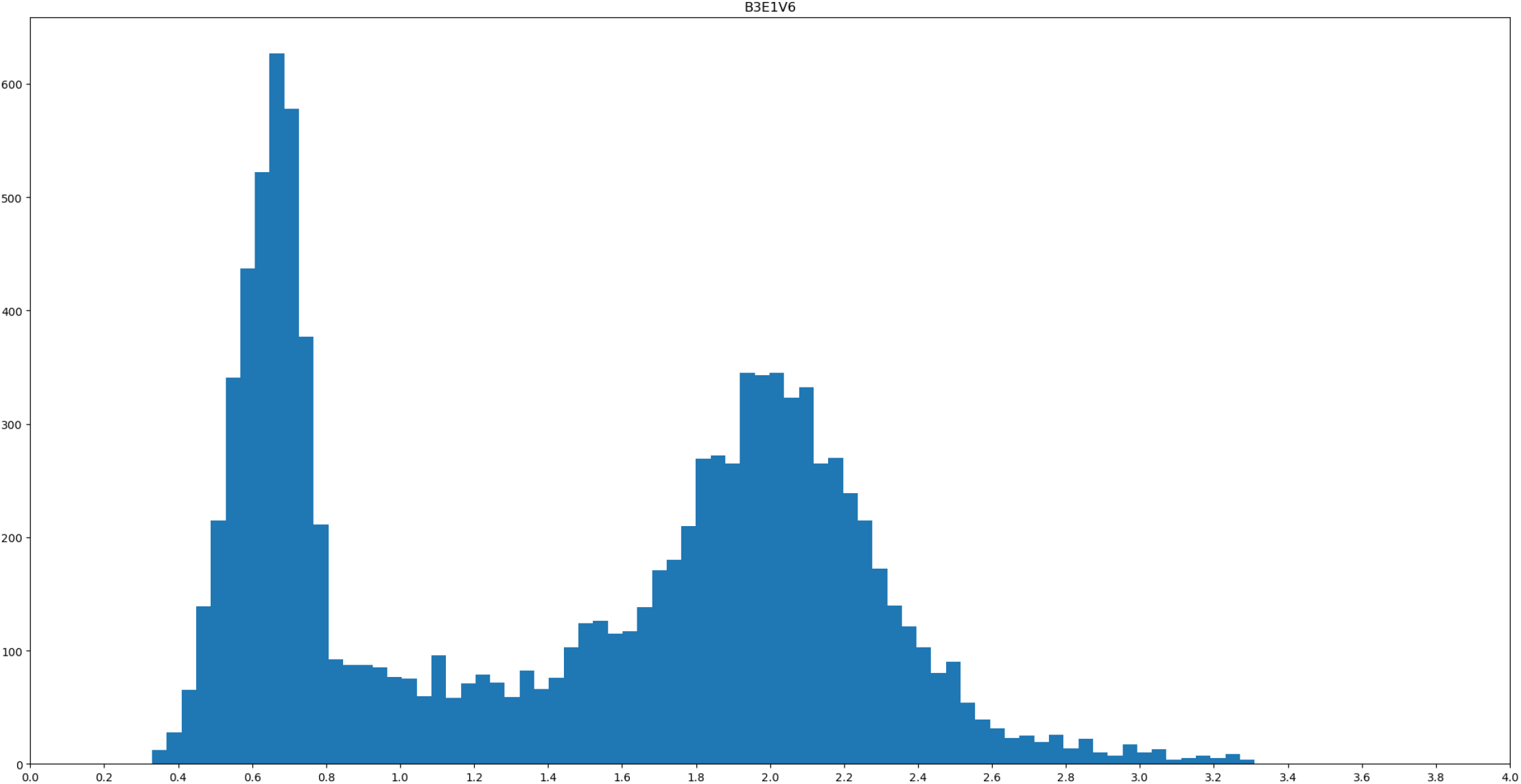
Sequence logo of profile B3E1V6 – neighborhood of the conserved histidine residue

**Figure S18:**
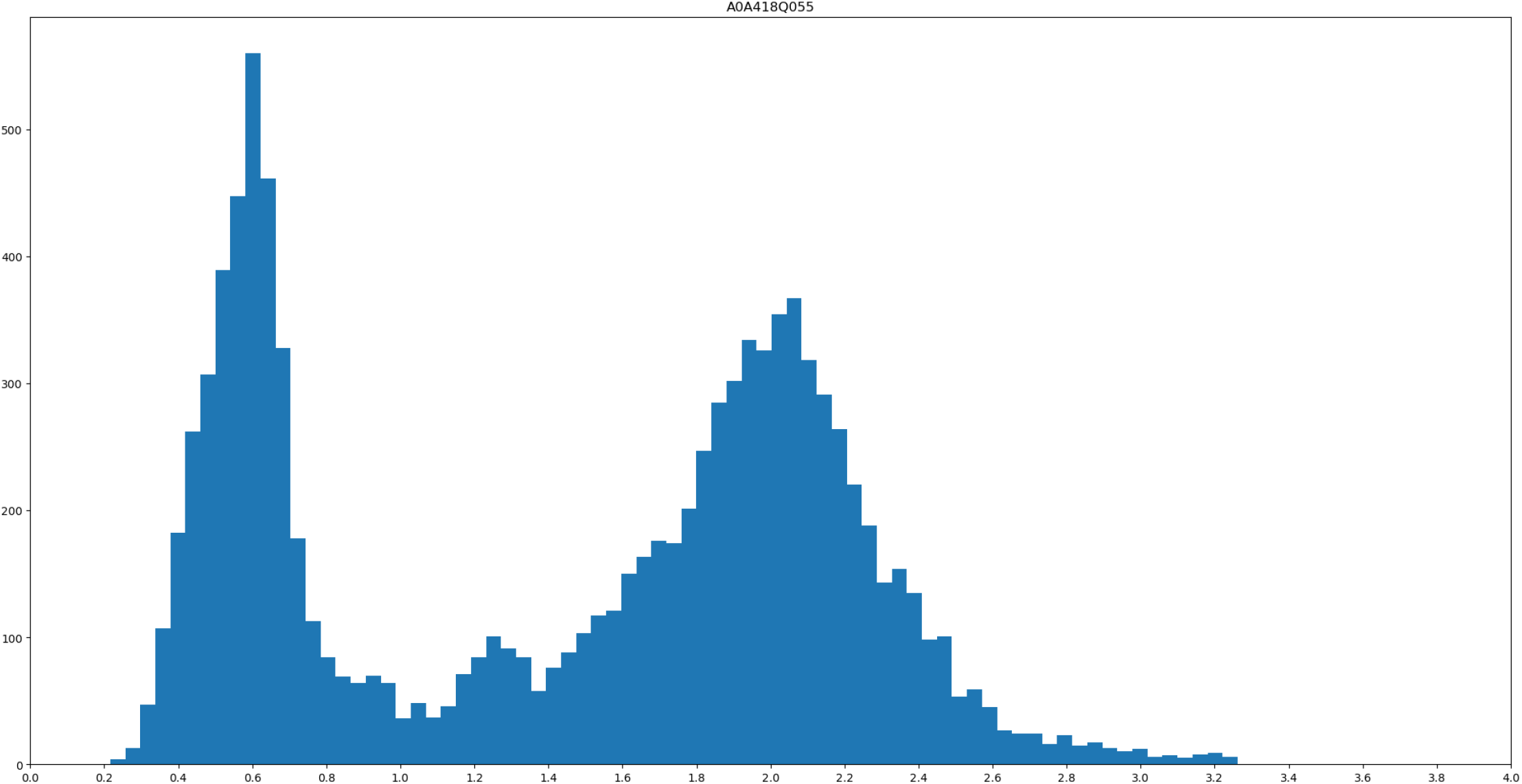
Sequence logo of profile A0A418Q055 – neighborhood of the conserved histidine residue (−5/+19).

**Figure S19:**
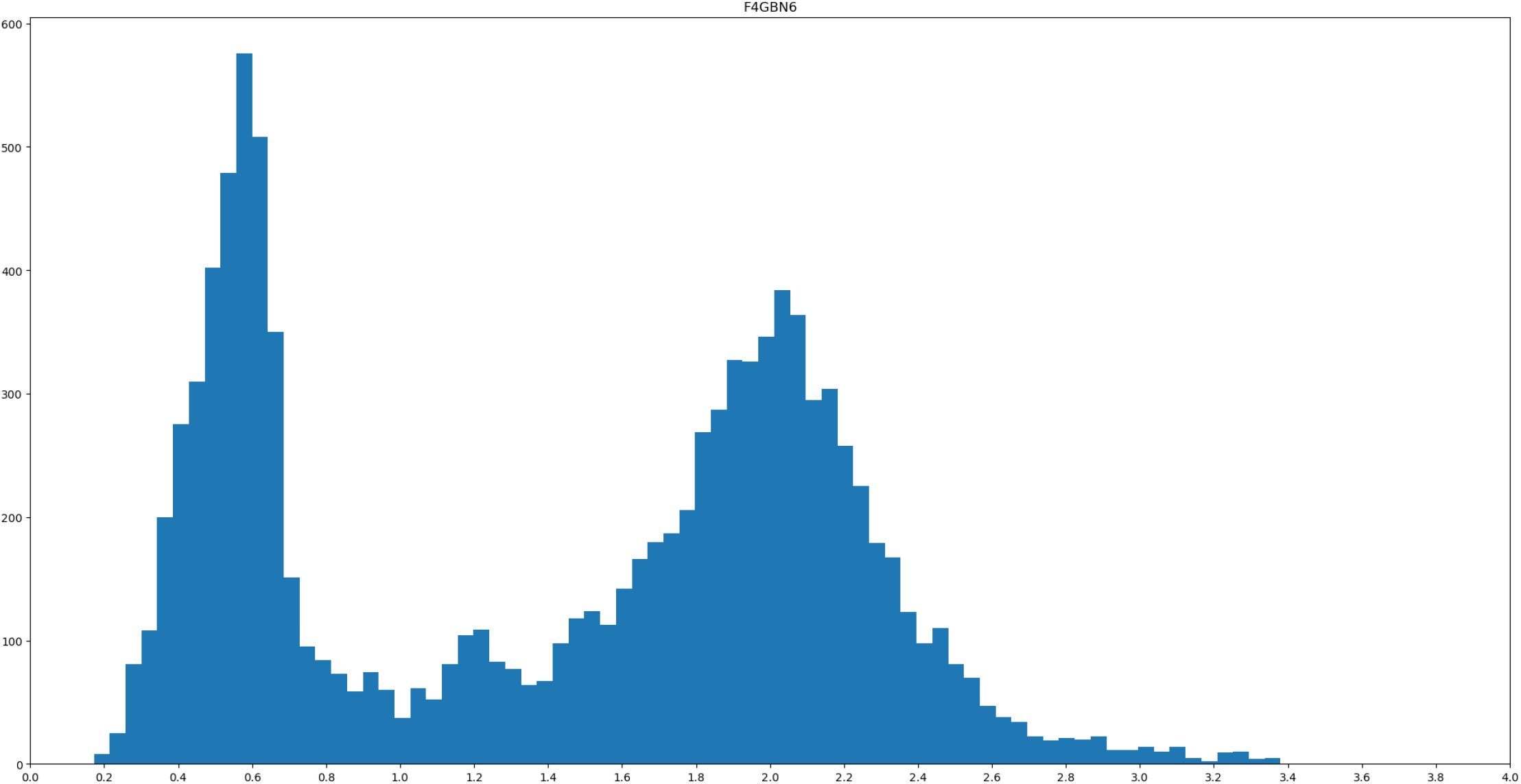
Sequence logo of profile F4GBN6 – neighborhood of the conserved histidine residue (−5/+19).

**Figure S20:**
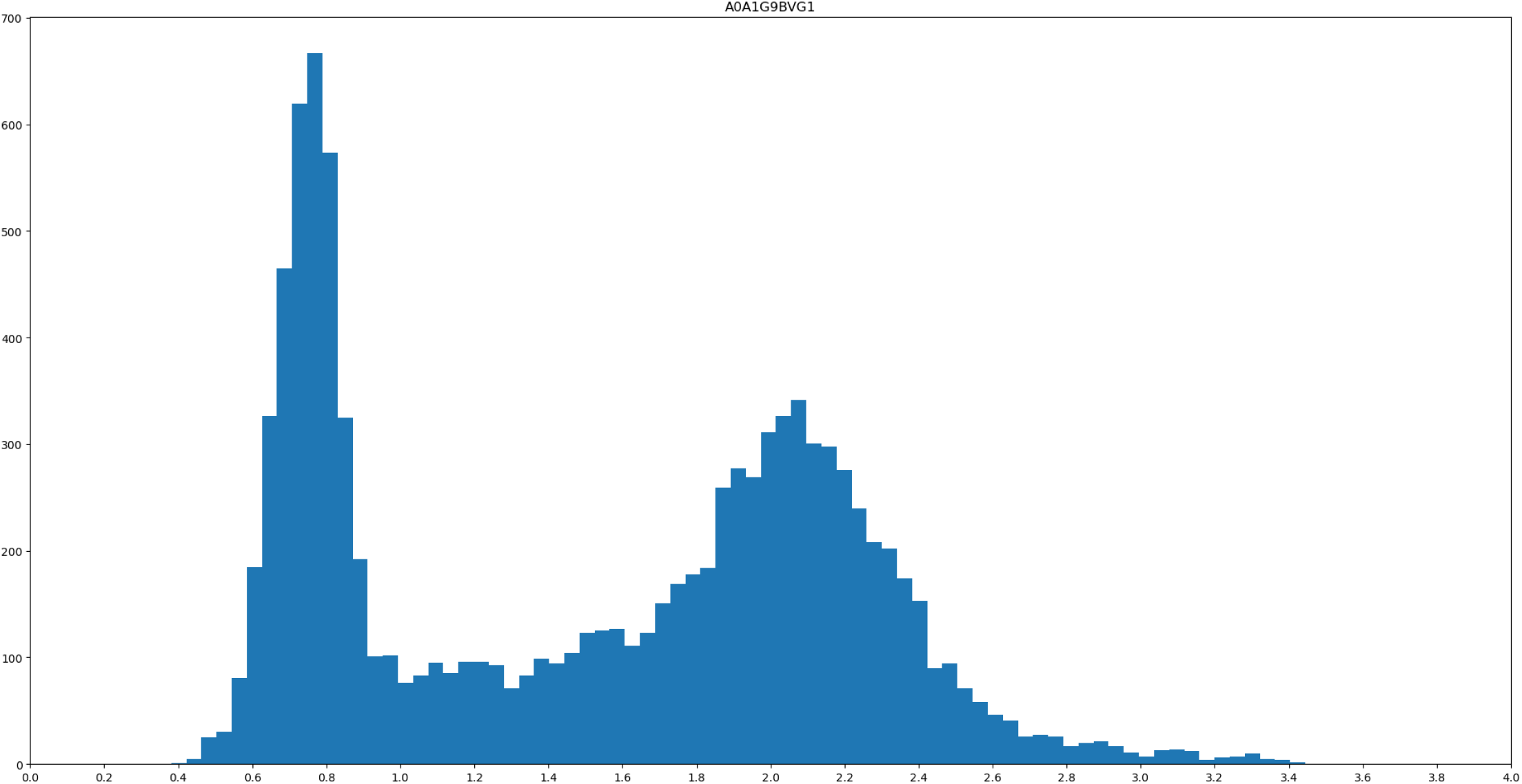
Sequence logo of profile A0A1G9BVG1 – neighborhood of the conserved histidine residue (−5/+19).

**Figure S21:**
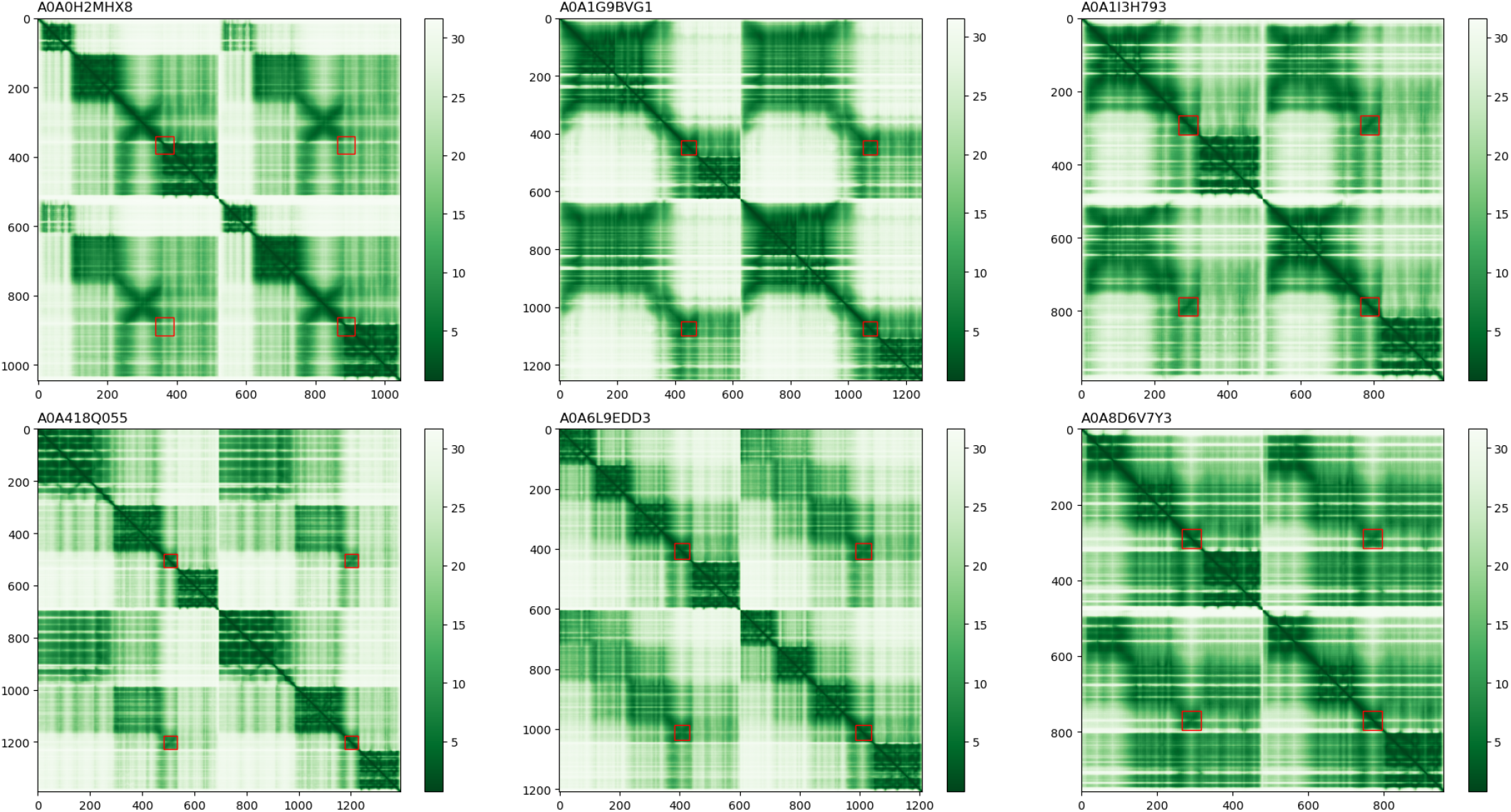
Interface PAE for six of the eighteen dimer models. Red rectangles highlight the regions aligning with the second version of the HisKA-like profiles, when 45 aa are considered after the conserved histidine residue (instead of 60).

**Figure S22:**
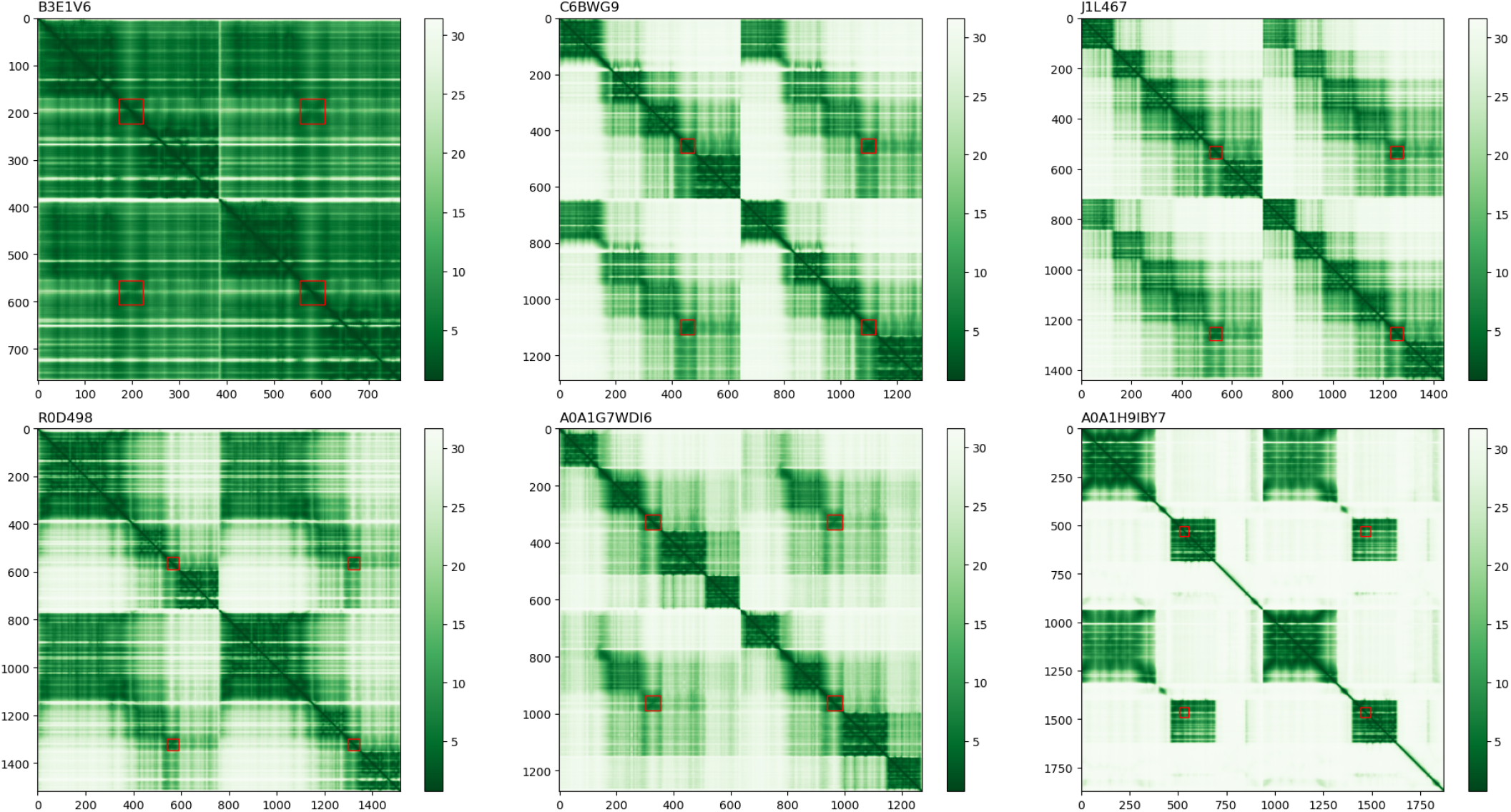
Interface PAE for six of the eighteen dimer models. Red rectangles highlight the regions aligning with the second version of the HisKA-like profiles, when 45 aa are considered after the conserved histidine residue (instead of 60).

**Figure S23:**
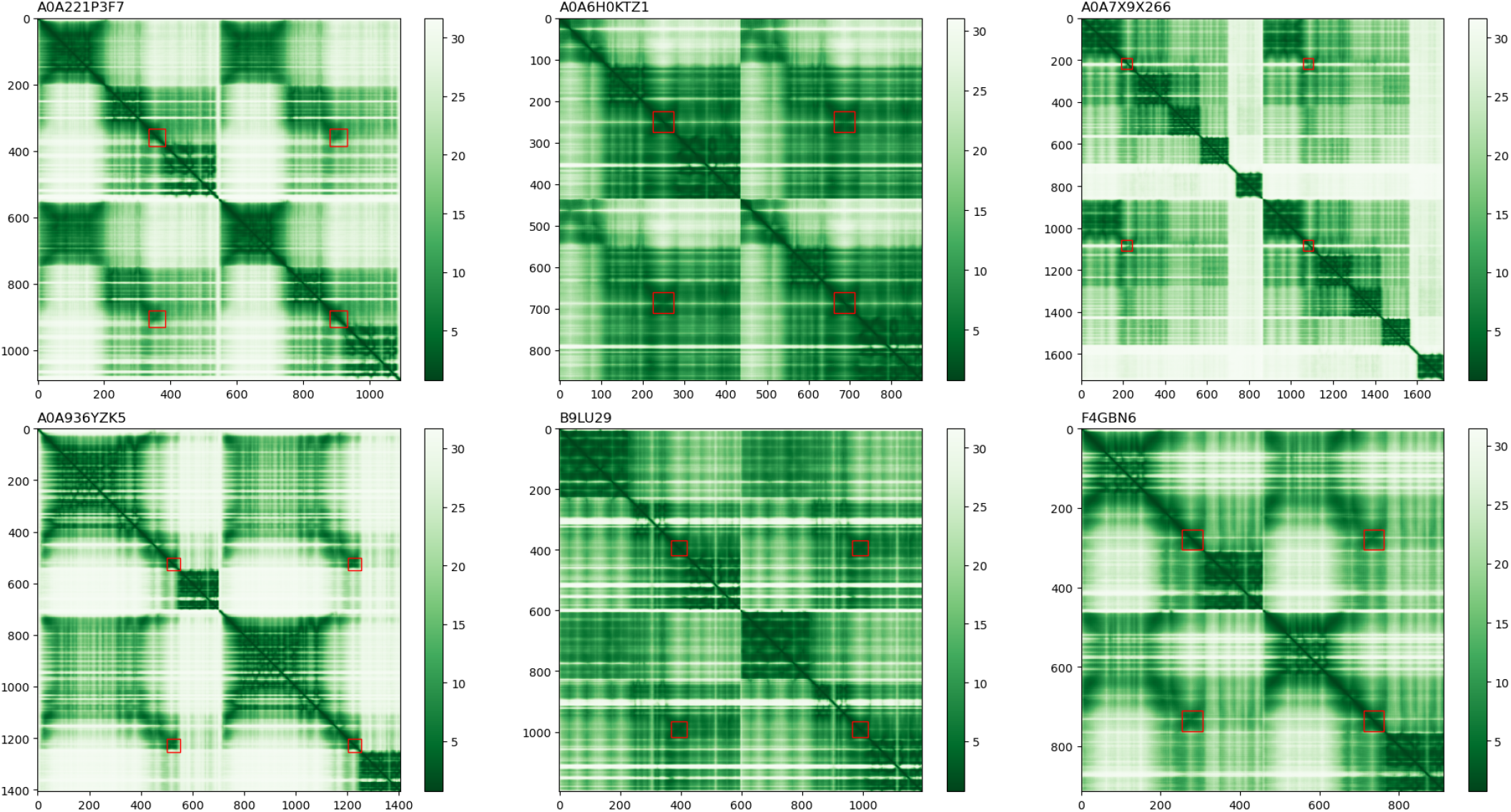
Interface PAE for six of the eighteen dimer models. Red rectangles highlight the regions aligning with the second version of the HisKA-like profiles, when 45 aa are considered after the conserved histidine residue (instead of 60).

